# Increased Th1 bias in memory T cells corresponds with protection from reinfection in *Plasmodium* infection, and is regulated by T cell-intrinsic STAT3

**DOI:** 10.1101/724963

**Authors:** Victor H. Carpio, Florentin Aussenac, Kyle D. Wilson, Alejandro V. Villarino, Alexander L. Dent, Robin Stephens

## Abstract

Hybrid Th1/Tfh cells (IFN-γ^+^IL-21^+^CXCR5^+^) predominate in response to persistent infections; however, molecular regulation of their function is poorly defined. In infection with *Plasmodium spp*, an IFN-γ^+^ T helper-1 (Th1) response controls initial parasitemia, while antibody and IL-21^+^CXCR5^+^ T follicular helper (Tfh) function effect final clearance. Here, we found that CD4-intrinsic Bcl6, Blimp-1 and STAT3 all regulate T-bet expression, which controls IFN-γ expression. While Bcl6 and Blimp-1 regulate the level of CXCR5, only T-bet and STAT3 affected the functional bias of the Th1/Tfh phenotype. Infected mice with STAT3-deficient T cells produced less antibody, and more IFN-γ^+^IL-21^−^CXCR5^lo^ T cells, significantly increasing protection from re-infection. Conversely, reduced Th1 bias in re-infected T-bet KO was reflected in prolonged secondary parasitemia. In summary, each feature of hybrid Th1/Tfh population in *Plasmodium* infection is uniquely regulated and the cytokine bias of memory T cells can be modified to enhance the effectiveness of the response.

## Introduction

Both cellular and humoral responses are essential for immunity from *Plasmodium* infection. In humans, CD4 T cells that produce IFN-γ in response to *P. falciparum* antigens accumulate with exposure, as do variant-specific antibodies, correlating with lower incidence of both parasitemia and hospitalization. A favorable ratio of IL-10 to TNF correlates with resistance from pathology in both mice and people (Li et al., 2003; Luty et al., 1999; May et al., 2000), and CD4 T cells protect immunodeficient mice from dying of *P. chabaudi* infection (Stephens et al., 2005). Both T helper-type 1 (Th1) promoting cytokines IL-12 and IFN-γ contribute to reduction of peak parasitemia by promoting parasite phagocytosis and generation of Th1-driven antibody isotypes (Stephens et al., 2005; Su and Stevenson, 2000; Xu et al., 2000). While IFN-γ production by T cells in response to *P. chabaudi* infection is initially strong, it becomes downregulated as infection becomes controlled. Thereafter, a much reduced but recrudescent parasitemia is cleared by germinal center (GC) derived antibody (Perez-Mazliah et al., 2017; Stephens et al., 2005). IL-21, made predominantly by CXCR5^+^ T cells, including T follicular helper (Tfh), is required for antibody isotype class switch, and contributes significantly to full clearance (Carpio et al., 2015; Perez-Mazliah and Langhorne, 2014; Perez-Mazliah et al., 2017).

In *P. chabaudi* infection, we and others have shown that many cells express both IFN-γ and IL-21 (Carpio et al., 2015; Perez-Mazliah et al., 2015). IFN-γ^+^IL-21^+^ CD4 T cells also occur in chronic LCMV, tuberculosis, and *Listeria* infections (Elsaesser et al., 2009; Li et al., 2016; Tubo et al., 2013). *In vitro*, prolonged TCR signaling and IL-12 drive T cells from the Th1 to the Tfh phenotype (Fahey et al., 2011; Schulz et al., 2009; Tubo and Jenkins, 2014). CXCR5^int^ effector T cells (Teff) have been demonstrated in other *Plasmodium* infections, and are capable of generating CXCR5^hi^PD-1^hi^ GC Tfh cells, therefore, are termed pre-Tfh (Ryg-Cornejo et al., 2016). We showed that the IFN-γ^+^IL-21^+^CXCR5^+^ T cells in *P. chabaudi* infection express the Tfh markers ICOS and BTLA, along with the IFN-γ-induced chemokine receptor CXCR3, and the primary transcription factors of both Th1 and Tfh (T-bet and Bcl6) (Carpio et al., 2015). This data led us to the term “hybrid Th1/Tfh” to describe any IFN-γ^+^ CD4 T cell also expressing IL-21 and/or CXCR5, functional markers of Tfh. Strikingly, IFN-γ^+^IL-21^+^ T cells are also the main source of IL-10 (Carpio et al., 2015; Perez-Mazliah et al., 2015), a critical cytokine as it prevents lethal pathology in *P. chabaudi* infected mice (Freitas do Rosario et al., 2012) and promotes antibody responses (Guthmiller et al., 2017). Th1/Tfh cells also preferentially expand during *P. falciparum* infection, where they have been termed Th1-like Tfh (Obeng-Adjei et al., 2015). However, mice with Bcl6-deficient T cells generated both CXCR5^int^ and IFN-γ^+^IL-21^+^ T cells in *P. chabaudi* infection (Carpio et al., 2015), suggesting that these hybrid phenotype T cells are not defined by this essential Tfh transcription factor. Nevertheless, both Tfh and Th1/Tfh cells help B cells make *Plasmodium*-specific antibodies, though Th1/Tfh (defined by CXCR3^+^, Ly6C^+^ or NK1.1^+^ in addition to CXCR5) help less efficiently as measured *in vitro* (Obeng-Adjei et al., 2015; Wikenheiser et al., 2018; Zander et al., 2017). This is likely due to an antagonism regulating Tfh effector functions through IFN-γ or TNF and T-bet expression, which inhibit GC Tfh, GC B cell formation, and IgG production in response to infection with *P. berghei* ANKA (Ryg-Cornejo et al., 2016). Therefore, the Th1/Tfh hybrid T cell population producing IFN-γ, TNF, IL-21 and IL-10 are likely to concurrently provide cellular protection and limit the large humoral response, which has been called hypergammaglobulinemia. It is not well understood which differentiation pathways control these effector cytokines, particularly in persistent infections. Therefore, we have investigated the molecular regulation of T cell cytokine production and phenotype in response to infection with *Plasmodium spp*.

Classically, committed IFN-γ^+^ Th1 cells are generated by antigen stimulation in the presence of IL-12, which signals through STAT4 (Hsieh et al., 1993) and increases levels of the master Th1 regulator, T-bet (Szabo et al., 2000). Th1 cells express CXCR3, but not CXCR5, which allows them to migrate away from the B cell follicle into the red pulp and inflamed tissues. The Tfh cell lineage-determinant transcription factor is Bcl6 (Johnston et al., 2009; Nurieva et al., 2009). Fully differentiated GC Tfh cells are identified as CXCR5^hi^PD-1^hi^ and their generation depends on the Tfh cell lineage-determinant transcription factor Bcl6 (Johnston et al., 2009; Nurieva et al., 2009). Many cytokines that regulate Tfh development, including IL-6, IL-27 and IL-21, signal through STAT3 (Crotty, 2014). IL-6 signaling through STAT3 secures Tfh programming by limiting Th1 differentiation (Choi et al., 2013). IL-27 signaling through STAT3 induces IL-21 production in T cells (Batten et al., 2010), which in turn promote Tfh development (Nurieva et al., 2008). *In vitro* and in response to viral infection, STAT3-deficient T cells have a defect in Tfh differentiation (Ray et al., 2014). Humans with dominant-negative mutations in STAT3 show compromised Tfh development (Ma et al., 2012a). However, over the last few years, several lines of evidence suggest a complex regulation of Th1 and Tfh where lineage determination is intertwined at the molecular level (Weinmann, 2014). For example, the transcription factor Blimp-1 can inhibit both Tfh and Th1 differentiation via transcriptional inhibition of Bcl6 and T-bet, respectively (Cimmino et al., 2008; Johnston et al., 2009). In the context of persistent infection, Blimp-1 also controls IL-10 production by Th1 cells (Parish et al., 2014).

Both Th1 and Tfh responses are critical for malaria immunity; however, the ideal balance between these T cell subsets remains unclear. Therefore, we investigated the roles of STAT3, T-bet, Bcl6 and Blimp-1 in the development of hybrid Th1/Tfh cells during persistent *P. chabaudi* infection in order to identify protective responses. We found that in contrast to the Th1/Tfh hybrid cells found in wildtype (WT) mice upon infection, T cells from T cell-specific STAT3 deficient mice (STAT3 TKO, Stat3^fl/fl^CD4^Cre^) preferentially differentiated into Th1 memory cells (IFN-γ^+^IL-21^−^T-bet^hi^). Strikingly, STAT3 TKO mice were 100% protected from reinfection, while T-bet deficient mice had no Th1 memory cells and higher parasitemia. Both mice had reduced serum levels of *Plasmodium*-specific IgG2b, the Th1 isotype, suggesting that the strong positive effect on parasitemia in STAT3 TKO mice was due to improved Th1 memory. Mechanistically, T-bet and not STAT1 or STAT4 regulated IFN-γ production by T cells, and T cell intrinsic expression of STAT3, Bcl6 and Blimp-1 each regulated T-bet expression during the peak of infection. Therefore, STAT3 is a key player regulating the cytokine plasticity of memory T cells in malaria. This data supports the hypothesis that Th cell pluripotency allows continued responsiveness promoting control of persistent infections and host homeostasis.

## RESULTS

### *Plasmodium* infections induce hybrid Th1/Tfh and GC Tfh cells

We have previously reported the presence of hybrid Th1/Tfh expressing both Tfh markers (CXCR5, ICOS, BTLA, IL-21 and Bcl6) and Th1 markers (CXCR3, IFN-γ, T-bet), as well as the regulatory cytokine IL-10 in *P. chabaudi* infection on day 7 p.i. (Carpio et al., 2015). To document the kinetics of the Th1 and Tfh marker phenotypes of Teff during blood-stage rodent *Plasmodium* infection, we infected C57BL/6J mice with *P. chabaudi* (AS) or *P. yoelii* (17XNL) infected red blood cells (iRBCs, Figure S1A), and measured parasitemia, GC B cell numbers and the expression of Th1 and Tfh markers over the course of infection. GC B cells make a late appearance in the third week in response to both infections. Unbiased t-Distributed Stochastic Neighbor Embedding (t-SNE) analysis gated on CD4^+^ T cells (Figure S1B) shows the small cluster of GC Tfh (Bcl6^hi^CXCR5^hi^PD-1^hi^), and the larger islands of hybrid Th1/Tfh cells (IFN-γ^+^IL-21^+^CXCR5^+^) generated in response to both infections.

We identify Teff as CD44^hi^CD127^−^, as IL-7Rα (CD127) is transiently downregulated upon activation, with its expression on the fewest T cells at day 9 p.i. (Stephens and Langhorne, 2010). CD127 downregulation correlates with CD11a expression (Figure S1C), which is upregulated by TCR, but not cytokine, stimulation (McDermott and Varga, 2011), making CD127^−^ a marker of antigen specificity. In the first week of infection, there is an unusual increase of CXCR5 expression on Teff in response to both *P. chabaudi* and *P. yoelii*, and the CXCR5^+^ Teff contained the majority of IFN-γ^+^ cells (Figures 1A, S1D). GC Tfh cells are present in stable numbers starting in the first week in both infections, as previously suggested (Wikenheiser et al., 2016). Many Teff produce both IFN-γ and IL-21, averaging 41.67% of Teff in *P. chabaudi* and 48.57% in *P. yoelii* in the first week of infection (Figures 1B and S1E). The IFN-γ^+^IL-21^+^ Teff population decreases by half in the second week and then remains a stable presence. Boolean gating analysis using IFN-γ, IL-21, CXCR5, T-bet and Bcl6 at day 7 p.i. showed that 66.43% of Teff from *P. chabaudi* infected mice co-express IFN-γ^+^ and at least one marker of Tfh (IL-21 or CXCR5), while the IFN-γ^+^T-bet^+^ Th1 cells represent a modest fraction (2.91%) of the response (Figures 1C, S1F).

**Figure 1.**
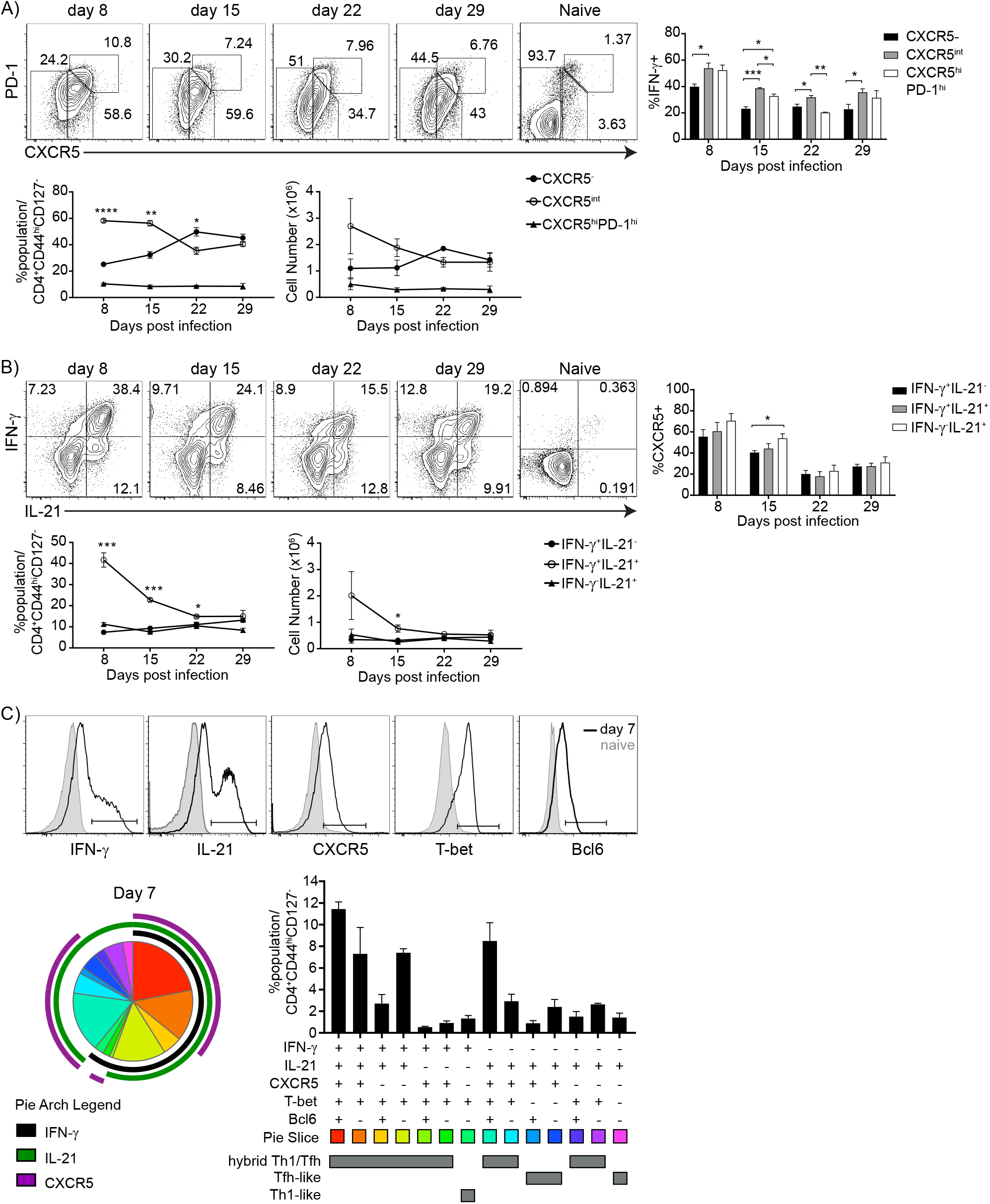
T helper differentiation during *P. chabaudi* infection resembles a hybrid Th1/Tfh phenotype. C57BL/6J mice were infected and splenocytes were analyzed at the indicated days.**(A)** Contour plots show expression of PD-1 and CXCR5 in CD4 Teff (CD44^hi^CD127^−^) and Naive (CD44^lo^CD127^+^). Below, line graphs show percentage (left) and numbers (right) of CXCR5^−^ (black filled dots), CXCR5^int^ (open circles) and CXCR5^hi^PD-1^hi^ (black filled triangles) populations. Right, bar graph shows percentage of IFN-γ^+^ in the different Teff subsets. **(B)** Contour plots show expression of IFN-γ and IL-21 in CD4 Teff and Naive. Below, line graphs show percentage (left) and numbers (right) of IFN-γ^+^IL-21^−^ (black filled dots), IFN-γ^+^IL-21^+^ (open circles), and IFN-γ^−^IL-21^+^ (black filled triangles) populations. Right, bar graph shows percentage of CXCR5^+^ in the different cytokine subsets. **(C)** Boolean gating of IFN-γ, IL-21, CXCR5, T-bet, and Bcl6 expression of CD4 Teff on day 7 p.i. Top, histograms show expression of IFN-γ, IL-21, CXCR5, T-bet, and Bcl6 in CD4 Teff. Bottom left, pie charts show the distribution of all the subsets. Bottom right, bar graph shows the percentage of the subsets. Data representative of 3 experiments with 3 mice/group.

In summary, *Plasmodium* infection primarily generates IFN-γ^+^IL-21^+^CXCR5^+^ hybrid Th1/Tfh effector cells, after a week of infection. Yet, GC do not form until week 2-3 of infection which coincides with a decrease in the proportions of IFN-γ^+^IL-21^+^CXCR5^+^ T cells. This finding is consistent with the observation that high production of IFN-γ or TNF after one week of infection inhibits GC formation (Hansen et al., 2017), and led us to question what features of infection regulate this unusual hybrid cytokine production by the dominant IFN-γ^+^IL-21^+^CXCR5^+^ T cells.

### A shorter infection results in fewer hybrid Th1/Tfh cells

Hybrid Th1/Tfh cells have been documented in human malaria patients (Obeng-Adjei et al., 2015) and in other persistent infections (Crawford et al., 2014; Nakayamada et al., 2011) using various combinations of Th1 and Tfh markers. However, acute infections promote independent differentiation of Th1 and Tfh populations (Curtis et al., 2010; Hale et al., 2013). Therefore, we hypothesized that a shorter *Plasmodium* infection would induce an altered T cell cytokine profile. We have previously shown that complete parasite clearance by the antimalarial drug Mefloquine (MQ) started on day 3 shaped the later Tcm/Tem ratio (Opata et al., 2015). As no qualitative change in phenotype was observed when drug treatment began on day 5 or 30 post infection (p.i.), there seems to be a limited window for determining the quality of T cell priming. Therefore, we tested if limiting the duration of infection by drug treatment would affect the generation of Th1/Tfh cells (Figure 2). MQ treatment of P. chabaudi-infected animals starting on day 3 cleared infection almost completely by day 5 (Figure 2A). MQ treatment has no known effect on immune cells at this low of a dose (Paivandy et al., 2014). Stopping the infection early (+MQ) decreased the numbers of Teff (Figure 2B), as expected due to the reduction in overall parasite present. There were also striking qualitative changes. Treatment of infection significantly reduced the proportions of CXCR5^int^ cells within the Teff population at day 7 p.i. (Figure 2C). Moreover, MQ-treated animals had a higher fraction of IFN-γ^+^IL-21^−^ Teff, and a strong reduction in the fraction and number of IFN-γ^+^IL-21^+^ T cells (Figure 2D). IFN-γ^+^IL-21^−^Teff from MQ-treated mice did not express CXCR5 (Figure 2E). Examining all markers together, MQ treatment reduced the proportion of hybrid IFN-γ^+^IL-21^+^CXCR5^+^ by 79.33 ± 3.41% and increased the proportions of IFN-γ^+^CXCR5^−^IL-21^−^ compared to untreated animals (Figure 2F), suggesting that this balance is affected by infection lasting longer than 3 days. GC B cell numbers were also increased at day 7 p.i. in treated compared to untreated mice (Figure 2G). In contrast, starting treatment on day 5, rather than day 3, had no effect on the fraction of IFN-γ^+^IL-21^+^ or CXCR5^int^ Teff (Figure S2). Therefore, we conclude that the stimuli responsible for priming of the hybrid Th1/Tfh cell phenotype, and leading to inhibition of GC B cell differentiation, occurs before day 5 of infection, the previously established window of initial priming in this infection (Opata et al., 2015).

**Figure 2.**
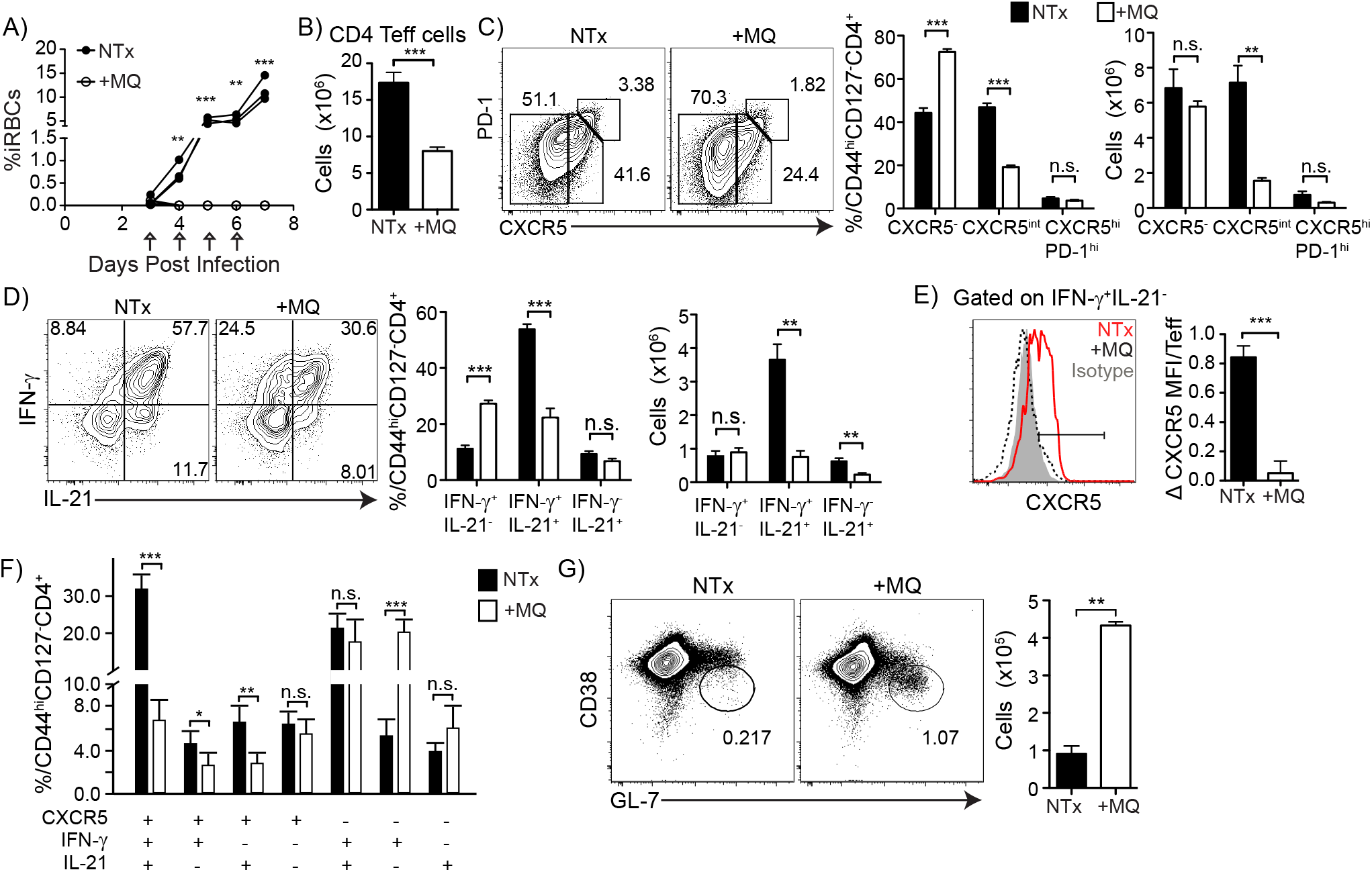
Drug-cured mice have fewer IFN-γ^+^IL-21^+^CXCR5^+^ hybrid Th1/Tfh, and more IFN-γ^+^IL-21^−^CXCR5^−^ Th1-like cells. C57BL/6J mice were infected, and one group was treated with mefloquine (MQ) starting day 3 and splenocytes were analyzed at day 7 p.i. **(A)** Parasitemia curve from not-treated (NTx, black filled circles) and treated (+MQ, open circles) groups. Arrows depict treatment days. **(B)** Bar graph shows CD4 Teff numbers in NTx (black) and +MQ (white) at day 7 p.i. **(C)** Contour plots show expression of PD-1 and CXCR5 and **(D)** IFN-γ and IL-21 in Teff. Bar graphs show percentages and numbers of Teff subsets. **(E)** Histogram shows CXCR5 expression in IFN-γ^+^IL-21^−^ Teff at day 7 p.i. Bar graph shows CXCR5 MFI (Mean Fluorescence Intensity) fold change over isotype control. **(F)** Boolean gating analysis of all possible combinations of CXCR5, IFN-γ and IL-21 expression by Teff. **(G)** Contour plots show expression of CD38 and GL-7 in B cells (B220^hi^MHC-II^hi^). Bar graph shows numbers of GC B cells. Data representative of 3 experiments with 3-4 mice/group.

### T-bet controls IFN-γ production

Basal levels of T-bet expression can be driven by TCR signaling, IFN-γ and STAT1. Higher levels of T-bet upregulate IL-12Rβ2, promoting IL-12 signaling through STAT4, to drive full Th1 commitment (Afkarian et al., 2002; Szabo et al., 2000). STAT4 (Figure S3A) was not required for the generation of IFN-γ^+^IL-21^+^ Teff, but it was critical for GC Tfh differentiation, in accordance with recent reports (Figure S3B (Weinstein et al., 2018)). T cell intrinsic STAT1 was also not definitively required for generation of IFN-γ^+^IL-21^+^ T cells (Figure S3C). It is of note, that we have not observed T-bet^hi^ Th1 committed cells in response to *P. chabaudi* infection in WT or even IL-10 KO mice (Carpio et al., 2015). Nevertheless, we investigated the role of T-bet in generation of hybrid Th1/Tfh cells. T-bet deficient (T-bet KO) mice infected with *P. chabaudi* had increased CXCR5^int^ Teff (Figure 3A), but a large decrease in the overall numbers of Teff compared to WT on day 7 p.i. (Figure 3B). Within the Teff population, there was a strong reduction in IFN-γ production in T-bet KO, and a strong increase in the fraction and number of IL-21^+^IFN-γ^−^ Teff (Figure 3C). Boolean gating analysis revealed a reduction in hybrid IFN-γ^+^IL-21^+^CXCR5^+^ cells, and a shift towards more Tfh-like Teff (IFN-γ^−^ IL-21^+^CXCR5^+/-^) in the absence of T-bet (Figure 3D). However, we found no significant increase in the number of GC B cells (Figure 3E). Supporting an important role for IFN-γ^+^ Teff in control of this infection, 40% of T-bet KO mice died from infection (Figure S3D). T-bet KO mice that survived the infection, did not control parasitemia as well as WT (Figure S3E), and had worse weight loss and hypothermia (Figure S3F). These data suggest that T-bet is required for IFN-γ production and control of parasitemia in *P. chabaudi* infection.

**Figure 3.**
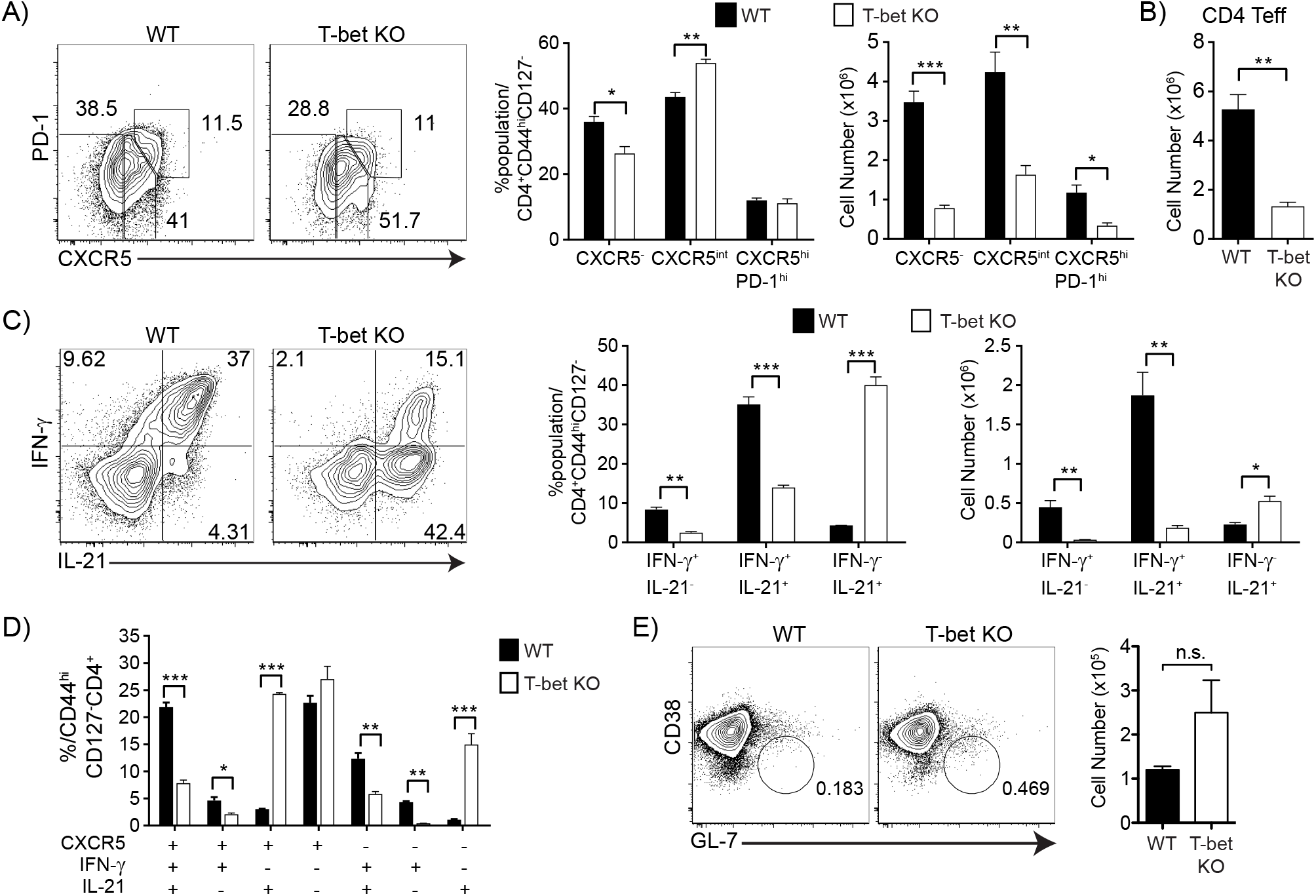
T-bet deficiency reduces hybrid Th1/Tfh and Th1 but promotes Tfh differentiation in *P. chabaudi* infection. T-bet KO and WT animals were infected and splenocytes were analyzed at day 8 p.i. Contour plots show expression of **(A)** PD-1 and CXCR5, and **(C)** IFN-γ and IL-21 in Teff. Bar graphs show percentages and numbers. **(B)** Bar graph shows CD4 Teff numbers in WT (black) and T-bet KO (white). **(D)** Boolean analysis of CXCR5, IFN-γ and IL-21 within WT (black) and T-bet KO (white) Teff. **(E)** CD38 and GL-7 in B cells (B220^hi^MHC-II^hi^). Bar graph shows numbers of GC B cells. Data representative of 2 experiments, 3-8 mice/group.

### Effector T cells deficient in STAT3 shift hybrid Th1/Tfh to a stronger Th1 bias

Because STAT3 promotes the Tfh phenotype (Batten et al., 2010; Choi et al., 2013; Nurieva et al., 2008; Ray et al., 2014), we hypothesized that STAT3 could also be a transcriptional regulator of the phenotype or function of hybrid Th1/Tfh cells. To test this hypothesis, we infected *Stat3*^fl/fl^CD4^Cre^ (STAT3 TKO) and *Stat3*^fl/fl^ (WT) animals with *P. chabaudi* or *P. yoelii* and analyzed splenocytes at day 7 or 10 p.i. by flow cytometry, respectively. We found no significant differences in the proportions of GC Tfh (Figure 4A and S4A). Infected STAT3 TKO mice did show a reduction in the fraction of IFN-γ^−^IL-21^+^ Teff (Figures 4B and S4B), supporting reports that STAT3 signaling promotes IL-21 expression. Infected STAT3 TKO mice also showed a decrease in the fraction of hybrid Th1/Tfh cells (IFN-γ^+^IL-21^+^CXCR5^+^), and an increase in the fraction of Th1-like cells (IFN-γ^+^IL-21^−^CXCR5^−^), particularly in *P. yoelii* infection (Figures 4C and S4C). The shift towards a stronger Th1 response in the absence of T cell STAT3 is evident from the increase in the CXCR5^int/lo^T-bet^hi^ population compared to WT (Figures 4D and S4D), suggesting a continuum (from Th1 to Tfh bias) rather than separate subsets. Although IL-6 (Figure S5A) and IL-27 (Figure S5B) signal through STAT3 and influence Tfh and Th1 differentiation respectively in other models, neutralizing these cytokines did not eliminate generation of IFN-γ^+^IL-21^+^ T cells. Generation of P. chabaudi-specific antibody was also affected by STAT3 deficiency in T cells. IgG titers were significantly less at day 35 p.i. in STAT3 TKO mice, while the relative concentration of IgM was not affected (Figure 4E). In addition, the proportion of GC B cells was significantly reduced in STAT3 TKO mice at days 20 and 55 p.i. (Figure 4F, d55 not shown). Parasitemia was prolonged (Figure S5C) and *P. chabaudi*-infected STAT3 TKO mice had a consistently prolonged bout of immunopathology as shown in the hypothermia, but not weight loss, data (Figure S5D). However, no STAT3 TKO mice died of *P. chabaudi* infection (n=40). The shift towards a stronger Th1 phenotype in STAT3 deficient animals indicates that STAT3 regulates the bias of hybrid Th1/Tfh cells during *Plasmodium* infection, though it also somewhat slows control of parasite increasing pathology in the first infection.

**Figure 4.**
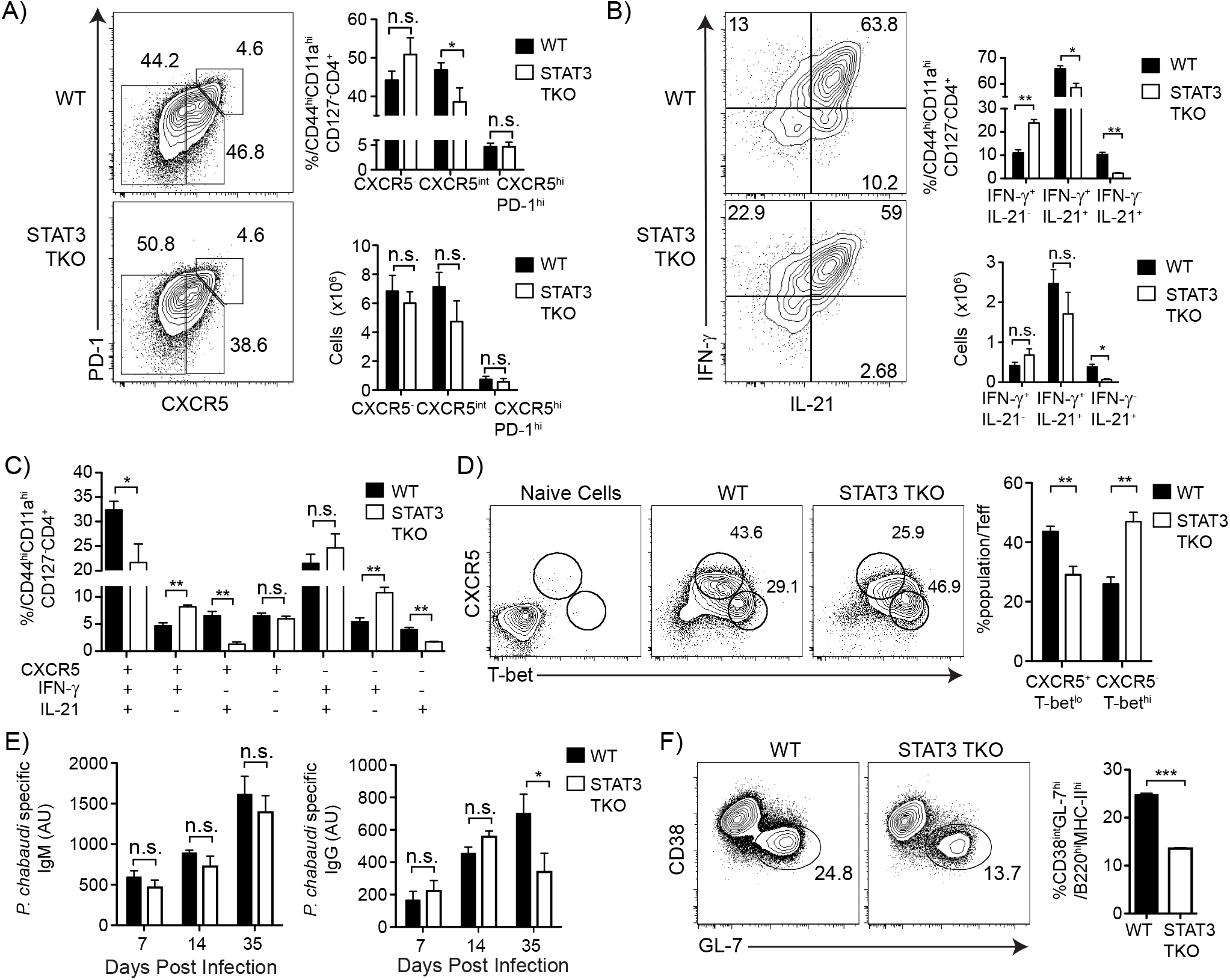
STAT3 deficiency reduces hybrid Th1/Tfh and Tfh but increases Th1 bias in *P. chabaudi* infection. *Stat3*^fl/fl^CD4^Cre^ (TKO) and *Stat3*^fl/fl^ (WT) animals were infected and splenocytes were analyzed at day 7 p.i. Contour plots show fraction of subsets using **(A)** PD-1 and CXCR5, and **(B)** IFN-γ and IL-21 gated on Teff. Bar graphs show percentages and numbers of subsets. **(C)** Boolean gating of CXCR5, IFN-γ, and IL-21 within WT (black bars) and STAT3 TKO (white bars) Teff. **(D)** Contour plots show expression of CXCR5 and T-bet in Teff. Bar graph shows percentages CXCR5^+^T-bet^lo^ and CXCR5^−^T-bet^hi^ subsets. **(E)** Bar graphs showing IgM (left graph) and IgG (right graph) antibodies specific for a lysate of *P. chabaudi*-infected red blood cells determined by ELISA. **(F)** Contour plots show expression of CD38 and GL-7 in B cells (B220^hi^MHC-II^hi^) day 20 p.i. Bar graph shows numbers. Data representative of 3 experiments, 3-4 mice/group.

### T-bet expression is regulated by Bcl6, Blimp-1 and STAT3

Bcl6 can bind and inhibit T-bet function (Oestreich et al., 2012), and indeed, we previously reported that Bcl6 levels correlate with the level of *Ifng* transcription in Teff in *P. chabaudi* (Carpio et al., 2015). Infected *Bcl6*^fl/fl^CD4^Cre^ (Bcl6 TKO) did not generate GC Tfh (Figure S6A). Nevertheless, Bcl6 deficient Teff could still acquire the CXCR5^+^IFN-γ^+^IL-21^+^ hybrid Th1/Tfh phenotype, though the level of CXCR5 was reduced. We also confirmed that there is an effect of T cell specific Bcl6 deficiency only during the clearance phase of parasitemia (Perez-Mazliah et al., 2017), and a slight increase in IL-10 (data not shown). Blimp-1 promotes terminal differentiation in plasma cells and CD8 Teff. However, in CD4 T cells, Blimp-1 can inhibit both T-bet and Bcl6, and is known to promote IL-10 production in *P. chabaudi* (Cimmino et al., 2008; Montes de Oca et al., 2016). Therefore, we tested Teff in infected *Prdm1*^fl/fl^CD4^Cre^ (Blimp-1 TKO) animals and found that the expression level of CXCR5 was increased, and that the relative fraction of CXCR5^+^IFN-γ^+^IL-21^+^ hybrid Th1/Tfh was increased (Figure S6B). Despite equal parasite levels, all of the Blimp-1 TKO died, likely due to elimination of IL-10 (Figure S6B). In summary, in *P. chabaudi* infection, Bcl6 and Blimp-1 primarily counter-regulate CXCR5 (and IL-10) expression levels, with smaller opposing effects on Th1-like IFN-γ^+^IL-21^−^CXCR5^−^ cells.

Based on the dramatic role of T-bet on IFN-γ production, we hypothesized that regulation of T-bet modulates the pathogenic potential of Th1-like cells *in vivo* and measured its expression in T cells in each infection. In WT animals, we previously showed that even though day 9 is the peak of IFN-γ^+^ Teff numbers, T-bet is downregulated from day 7 to day 9 p.i, suggesting a failure of Th1-commitment (Carpio et al., 2015). Here, in Bcl6 TKO animals, T-bet expression was maintained in Teff from day 7 to day 9 p.i, and Blimp-1 expression was increased at day 9 p.i. (Figure 5A). Blimp-1 deficient T cells showed an earlier effect on the expression of T-bet, which was already higher by day 7 p.i. (Figure 5B). As expected, Blimp-1 TKO Teff had an increase in Bcl6 at day 7 p.i. STAT3 TKO Teff also had more T-bet and Blimp-1 expression at day 7 p.i (Figure 5C), but Bcl6 expression was not affected, though it was reduced in *P. yoelii* infection of STAT3 TKO (Figure S4F). In conclusion, Bcl6, Blimp-1 and STAT3 are negative regulators working in concert to control the expression of T-bet, IFN-γ, CXCR5 and each other, in Teff during *Plasmodium* infection.

**Figure 5.**
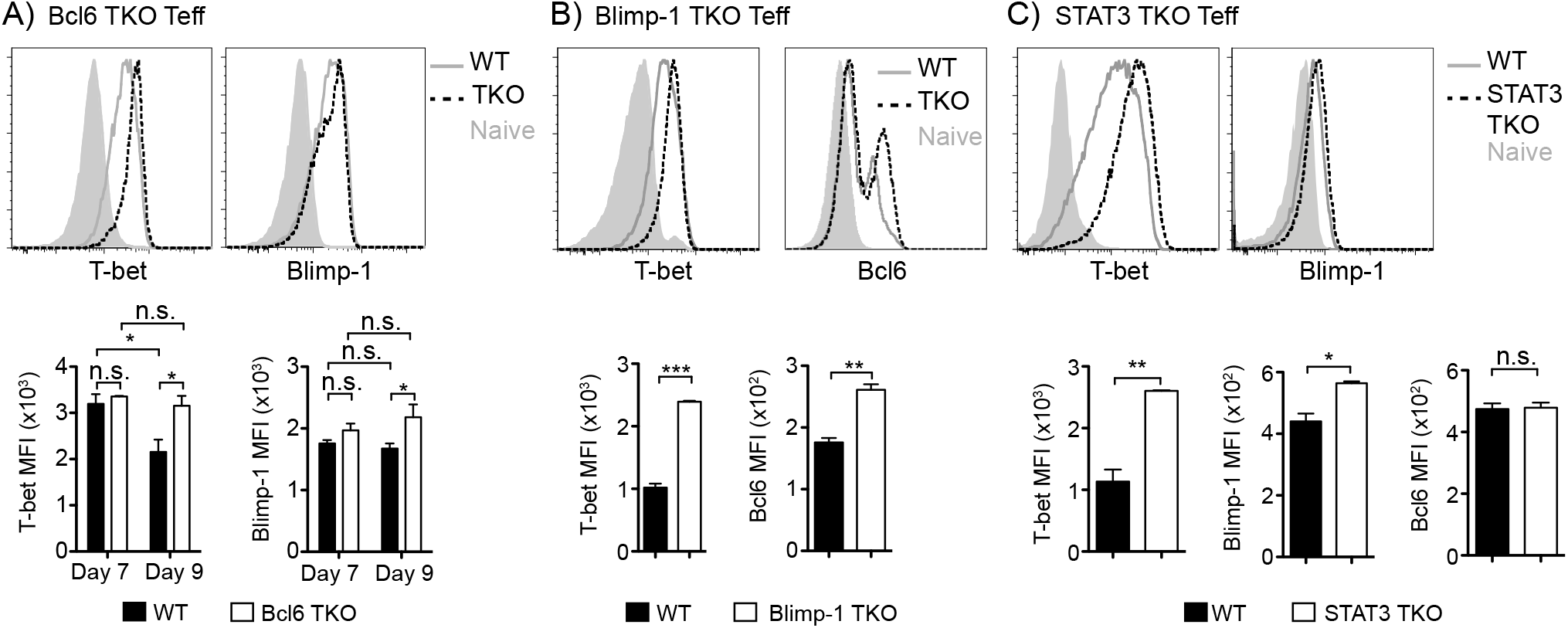
Bcl6, Blimp-1 and STAT3 control T-bet expression during *P. chabaudi* infection. TKO and WT animals were infected and splenocytes were analyzed at day 7 and 9 p.i. **(A)** Histograms showing T-bet (left) and Blimp-1 (right) expression in Teff from Bcl6 TKO (dotted line), WT (gray line), and naive (gray filled line) cells at day 9 p.i. Bar graphs shows average MFI of T-bet and Blimp-1 at days 7 and 9 p.i. **(B)** Histograms showing T-bet (left) and Bcl6 (right) expression in Teff from Blimp-1 TKO (dotted line) and WT (gray line) animals, and naive (gray filled line) cells at day 7 p.i. Bar graphs shows average MFI of T-bet and Bcl6 at day 7 p.i. **(C)** Histograms showing T-bet (left) and Blimp-1 (right) expression in Teff from STAT3 TKO (dotted line) and WT (gray line) animals, and naive (gray filled line) cells at day 7 p.i. Bar graphs shows average MFI of T-bet, Blimp-1, and Bcl6 staining at day 7 p.i. Data representative of 3 experiments with 3-4 mice/group.

As the hybrid Th1/Tfh phenotype is increased when infection lasts longer than 3 days, and there were significant differences between the roles of Bcl6, Blimp-1 and STAT3 previously reported and those described here, we next compared the effects of transcription factor deficiency on Teff in animals with MQ-treated infections. WT and TKO animals were infected, and one group of each was treated with MQ starting at day 3 p.i. (Figure S7). Data is displayed as contour plots and quantified as a ratio of TKO over WT to highlight the degree of the effect of removal of each transcription factor in the longer (NTx) or the shorter (+MQ) infection. Both IFN-γ^+^IL-21^−^ and hybrid IFN-γ^+^IL-21^+^ Teff were significantly increased in the short-term infection in Bcl6 TKO mice compared to WT, which supports *in vitro* data showing a large effect of Bcl6 on IFN-γ expression (Figure S7A, (Oestreich et al., 2011)). On the other hand, Blimp-1 seems to play a larger role in longer infection, as only *P. chabaudi*-infected Blimp-1 TKO (NTx) had fewer IFN-γ^+^IL-21^−^ with a concomitant increase in hybrid IFN-γ^+^IL-21^+^. This is consistent with its role in regulation of IL-10, a regulator of prolonged inflammation. On the other hand, STAT3 regulates IFN-γ in both long and short infections. This is clearly shown in the strong increase of IFN-γ^+^IL-21^−^ Teff in STAT3 TKO mice compared to WT. However, GC Tfh generation was only affected by STAT3 deficiency in the shortened infection, supporting the previously described role of STAT3 in Tfh in acute infection (Figure S7B, (Ray et al., 2014)). Together, these results suggest that the role of each transcription factor is dependent on the duration of strong priming, presumably due to differential expression of cytokines and transcription factors driven by the milieu.

### Increasing Th1 bias in memory T cells coincides with lower parasitemia in reinfection

Given the increase of Th1 cells and effective clearance of persistent parasite in STAT3 TKO mice, and the strong impact of type-1 cytokines on parasitemia in mice and humans (Luty et al., 1999; Stevenson et al., 1995), we re-infected STAT3 TKO animals to test for immunity (Figure 6). To ensure parasite clearance after the first infection in both STAT3 TKO and WT, we treated with the anti-malarial drug Chloroquine (CQ), which effectively eliminates low levels of *P. chabaudi* parasitemia (Hunt et al., 2004). STAT3 TKO mice controlled a strong second challenge (1 × 10^7^ iRBC) completely, with infection becoming undetectable by day 3 post reinfection (p.r.i, Figure 6A). WT mice showed significantly higher parasitemia that peaked around day 4, and was controlled by day 7 p.r.i. The proportion of IFN-γ^+^IL-21^−^ T cells was higher in STAT3 TKO mice at day 7 p.r.i, and the numbers of IFN-γ^+^IL-21^+^ Teff were less (Figure 6B). The numbers of both GC Tfh (Figure 6C) and GC B cells (Figure 6D) were significantly less in STAT3 TKO mice than WT at days 7 p.r.i. and day 55 p.i. (not shown). To determine if increased Th1 bias of Teff in the STAT3 TKO observed at the peak of the first infection was maintained into the memory phase, we analyzed antigen-experienced memory T cells (Tmem, CD11a^hi^CD49d^hi^CD44^hi^CD127^hi^) at day 55 p.i. Indeed, STAT3 deficient Tmem had more percentages of IFN-γ^+^IL-21^−^ cells (Figure 6E) and maintained a higher expression of T-bet (Figure 6F) than WT. Importantly, the levels of *P. chabaudi*-specific IgG and Th1-driven isotype, IgG2b, were significantly less in STAT3 TKO mice than WT (Figure 6G).

**Figure 6.**
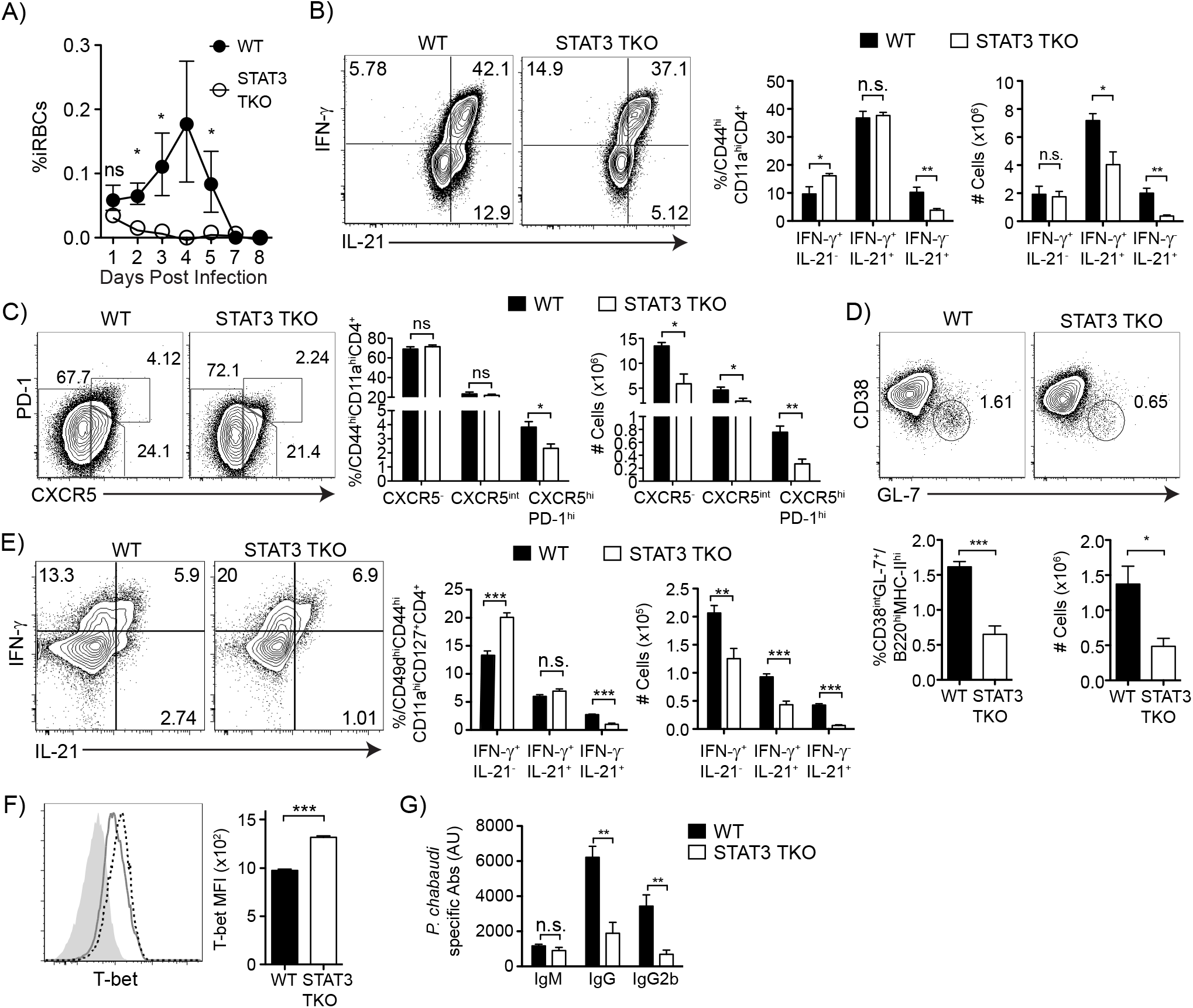
STAT3 TKO mice are protected from reinfection despite showing weaker humoral responses. STAT3 TKO and WT animals were infected. At day 60 p.i. both groups were treated with chloroquine (CQ) before reinfection. After 6 weeks mice were re-infected with 10^7^ iRBCs. **(A)** Parasitemia curve from WT (black filled circles) and STAT3 TKO (open circles) animals. Contour plots show expression of **(B)** IFN-γ and IL-21, and **(C)** PD-1 and CXCR5 by antigen-experienced CD4 T cells (CD44^hi^CD11a^hi^) from STAT3 TKO and WT animals at day 7-post reinfection. Bar graphs shows percentages and numbers. **(D)** Contour plots show expression of CD38 and GL-7 in B cells (B220^hi^MHC-II^hi^) at day 7-post reinfection. Below, bar graphs show percentages and numbers. **(E)** Contour plots show expression of IFN-γ and IL-21 in Tmem from STAT3 TKO and WT animals at day 55 p.i. (1^st^ infection). Bar graphs show percentages and numbers. **(F)** Histogram shows T-bet expression in Tmem from WT (gray line) and STAT3 TKO (dotted black line) animals, and naive (gray filled line) cells at day 55 p.i. (1^st^ infection). Bar graph shows MFI of T-bet. **(G)** Bar graphs showing IgM, IgG and IgG2b antibodies specific for a lysate of *P. chabaudi*-infected red blood cells determined by ELISA at day 7 post reinfection. Data representative of 2 experiments, 4-5 mice/group. Data in **(A)** is pooled from 2 independent experiments, 3-5 mice/group/experiment.

The increase in Th1-like (IFN-γ^+^IL-21^−^) T cells, decrease in *Plasmodium*-specific serum antibody, and concomitant very strong protection in STAT3 TKO mice supports a role for Th1 cells in reinfection. Therefore, we tested the importance of Th1 cells in immunity by giving T-bet KO mice a second infection (Figure 7). T-bet KO mice showed prolonged parasite growth compared to WT mice, with days 6 and 7 p.r.i. remaining uncontrolled (Figure 7A). This was the opposite phenotype to STAT3 TKO, as predicted. Upon *P. chabaudi* reinfection, Teff in T-bet deficient mice still produced IL-21, but little IFN-γ (Figure 7B). T-bet deficient mice had increased levels of *P. chabaudi*-specific IgG, but lower levels of Th1-isotype IgG2b (Figure 7C). In summary, T cell intrinsic STAT3 regulates the Th1 bias of memory T cells in *P. chabaudi* infection. Importantly, Th1 cells promote immunity, in addition to the role of pre-existing antibody, particularly IgG2b (Su and Stevenson, 2002).

**Figure 7.**
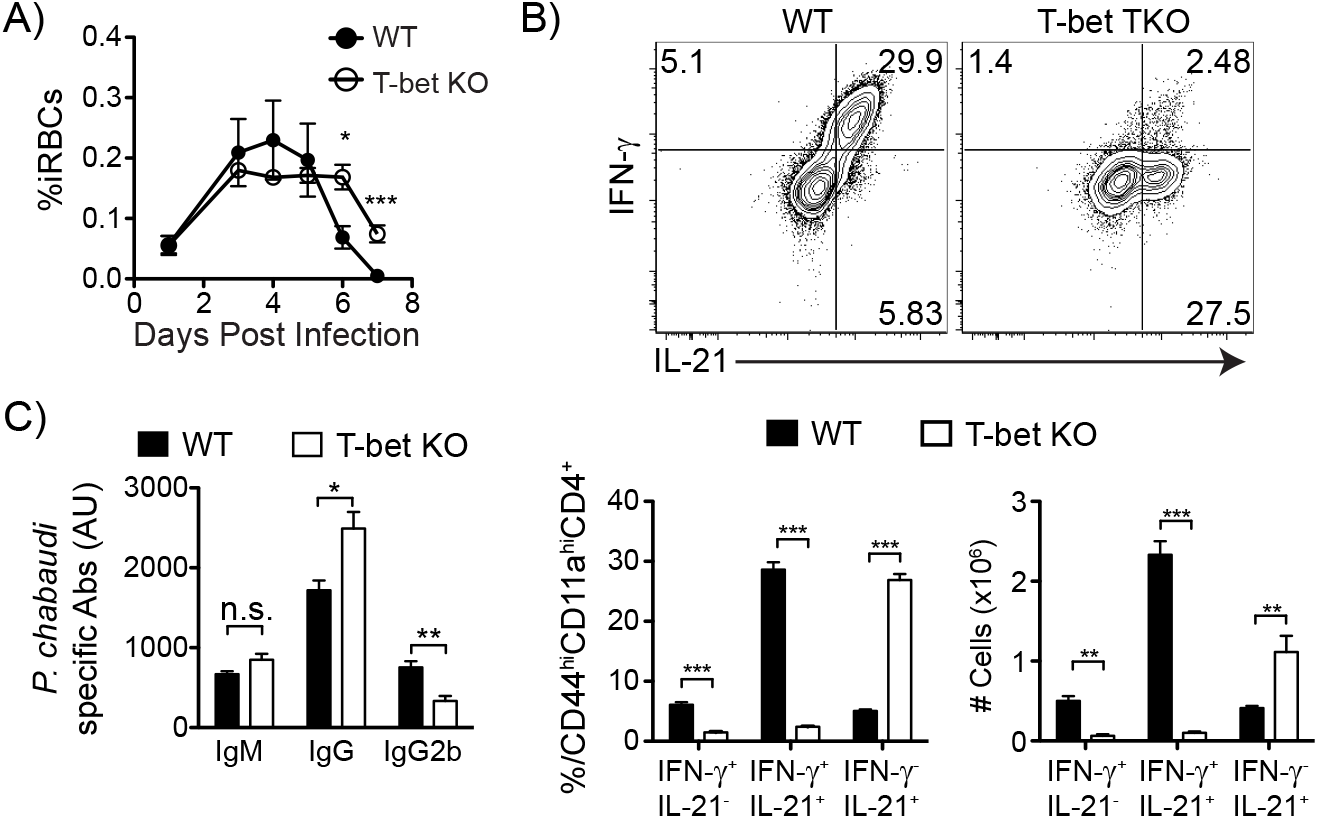
T-bet KO mice have no Th1 memory cells, less IgG2b antibodies, and do not control parasite after reinfection. T-bet KO and WT animals were infected. At day 60 p.i. both groups were treated with chloroquine (CQ) before reinfection. After 3 weeks mice were re-infected with 10^7^ iRBCs (A) Parasitemia curve from WT (black filled circles) and T-bet KO (open circles) animals. **(B)** Contour plots show expression of IFN-γ and IL-21 by antigen experienced CD4 T cells from T-bet KO and WT animals at day 7 post reinfection. Below, bar graphs show percentages and numbers. (C) Bar graphs showing IgM, IgG and IgG2b antibodies specific for a lysate of P. chabaudi-infected red blood cells determined by ELISA at day 7-post reinfection. Data representative of 1 experiment, 3-5 mice/group.

## Discussion

Both Th1 and Tfh cells are required to eliminate parasites in *Plasmodium* infection. Previous work on the immune response to *P. chabaudi* shows that IFN-γ controls the height of the peak of parasitemia, while Tfh and IL-21 are required for antibodies to eliminate the parasite (Amante and Good, 1997; Perez-Mazliah et al., 2015; Perez-Mazliah et al., 2017; Stephens et al., 2005; Su and Stevenson, 2002). We have found that both types of effector functions are combined in one cell type in this infection (Carpio et al., 2015). While there are certainly GC Tfh that make IFN-γ, we continue to term these multi-functional Teff cells found in persistent infections hybrid Th1/Tfh, rather than Th1-like Tfh, due to the active regulation of T-bet (which controls their IFN-γ and the small effect of Bcl6 (suggesting a more Th1-like regulation), as well as the lack of the true Tfh phenotype. It is important to note that in some staining combinations, two populations (i.e. CXCR5^int^T-bet^hi^, CXCR5^lo^T-bet^int^) appeared detectable within the hybrid population by FACS, and previously by scRNA-Seq (Lonnberg et al., 2017). We would argue that the CXCR5^int^ population we can detect here with improved CXCR5 staining, and the two plastic populations within it, do not represent truly differentiated populations, but actually two ends of a continuum. The fact that each feature of these cells is actively regulated by so many transcription factors highlights their plasticity, as a necessity of adapting to the current milieu. Therefore, we conclude that the Teff population in this infection is not made up of subsets, but is a plastic, heterogenous, hybrid population that is actively regulated by multiple inputs and transcription factors throughout the infection.

Protection from repeated episodes of malaria in humans correlates with serum IFN-γ and memory Th1 cells (Luty et al., 1999; Moormann et al., 2013; Stephens and Langhorne, 2010). CD4 T cells in adults from malaria endemic areas also express cytokines of multiple lineages including IFN-γ, IL-10 and IL-21, even in cells with a Tfh-like phenotype (Obeng-Adjei et al., 2015; Roetynck et al., 2013). While human Teff cells expressing both CXCR3 and CXCR5 can help B cells make antibody, mouse CXCR5^+^ Teff cells expressing markers of a high level of activation (Ly6C, NK1.1) are less effective helper cells *in vitro* than those expressing only CXCR5 (Obeng-Adjei et al., 2015; Zander et al., 2017). In humans, CXCR3^+^CXCR5^+^ T cells were not shown to correlate with *Plasmodium*-specific antibody levels (Obeng-Adjei et al., 2015). Strikingly, Th1/Tfh hybrid T cells (ICOS^+^CXCR3^+^CXCR5^+^) did correlate with antibody levels in influenza, where they were also shown to contain IFN-γ^+^IL-21^+^ cells (Bentebibel et al., 2013). Acute infections, like that caused by Listeria, can induce a stable Th1 memory phenotype (Curtis et al., 2010; Hale et al., 2013), while chronic LCMV and tuberculosis infections have T cell responses skewed away from a committed Th1 phenotype, and towards concomitant expression of Tfh markers in mice (Crawford et al., 2014; Li et al., 2016). Therefore, Th phenotype plasticity appears to be a shared feature of the immune response to persistent infections, and has been shown to be beneficial in control of tuberculosis (Khader et al., 2007; O’Shea and Paul, 2010). Similarly, our data suggest that preserving plasticity would be optimal for protection, i.e. the possibility that a subsequent infection drives Th1/Tfh hybrid cells towards Th1.

We found that T-bet drives much of the IFN-γ production in T cells in *P. chabaudi* infection. In addition, the expression of T-bet is highly regulated, including by STAT3, Blimp-1 and Bcl6, presumably to avoid immunopathology. T-bet expression kinetics also support ongoing regulation. We previously observed that T-bet is downregulated before day 9, the peak of both parasitemia and IFN-γ production (Carpio et al., 2015). Here, we show that this downregulation occurs in a Bcl6-dependent manner. While T-bet is generally induced by STAT1 in CD4 T cells (Afkarian et al., 2002) and upregulated upon IL-12 signaling by STAT4, neither STAT4 nor STAT1 deficiency negatively regulated the hybrid cytokine profile in T cells. This observation suggests that T-bet in this infection is induced primarily by TCR signaling and/or that there is some redundancy between STATs.

It seems that this hybrid T cell with a dual nature facilitates simultaneous cellular and humoral responses; however, the two types of responses clearly also regulate one another. T-bet in T cells has recently been shown to impair GC Tfh cell differentiation, which is essential for parasite elimination but not survival (Perez-Mazliah et al., 2017; Ryg-Cornejo et al., 2016). On the other hand, a recent study concluded that T-bet and STAT4 are actually required for GC Tfh development and GC formation during acute viral infection (Weinstein et al., 2018). We confirm that STAT4 is required for GC Tfh; however, we did not detect any change in the number of GC Tfh in T-bet deficient animals infected with P. chabaudi. T-bet, presumably in its capacity for driving IFN-γ in T cells and production of the IgG2 isotype of antibody, was essential for parasite control in both the first and second infections. Clearly, the regulation of T-bet and IFN-γ is a high priority for promoting an effective Teff response and survival of the animals given the multiple transcription factors involved. This regulation becomes particularly important when the innate response is not sufficient to limit infection by day 3. However, the myriad of cytokines produced in response to continued infection leads to plasticity due to the competition of effector differentiation programs within individual effector T cells.

Significant recent research has focused on the overlapping signaling cascades controlling the balance of Th1 and Tfh programs (Weinmann, 2014). IL-12, the primary cytokine responsible for induction of Th1 cells, can also induce IL-10 and can be essential for generation of Tfh *in vivo* (Saraiva et al., 2009; Weinstein et al., 2018). In addition, *in vitro* generated Th1 cells transiently express Bcl6 and IL-21 while Tfh transiently express T-bet (Fang et al., 2018; Nakayamada et al., 2011). T-bet can bind and inhibit the Tfh driving transcription factor, Bcl6 (Oestreich et al., 2012). While we previously showed that Bcl6 levels correlate with the level of *Ifng* transcription in these cells, the Bcl6 TKO do not have more IFN-γ^+^ T cells by intracellular cytokine staining here. In preliminary data, we did not see intermediate phenotypes in heterozygous mice, so we have not been able to test the role of reduced levels of each of the regulating transcription factors or their effects on parasitemia and the cytokine milieu. Both STAT3 and Bcl6 are reported to be required for Tfh differentiation, while Blimp-1 inhibits both Th1 and Tfh differentiation (Cimmino et al., 2008; Johnston et al., 2009; Ma et al., 2012; Nurieva et al., 2009; Ray et al., 2014). However, in our studies, deficiency in either Bcl6 or STAT4 in T cells eliminated GC Tfh, while STAT3 deficiency did not. STAT3 deficiency did however, reduce the proportions of GC Tfh in the setting of a shorter *P. chabaudi* infection, in agreement with previous reports that used acute infections as stimuli (Ray et al., 2014). In addition, we consistently observed that STAT3 deficient T cells did not develop into Tfh-like IFN-γ^−^IL-21^+^ Teff. The shorter infection also resulted in an increase of GC B cells, similar to the increase in GC B cells induced by inactivated *P. berghei* ANKA (Ryg-Cornejo et al., 2016). Bcl6 also reduced the level of expression of CXCR5, while Blimp-1 had the opposite effect. Our data suggest that Bcl6 and Blimp-1 have a stronger effect on regulation of CXCR5 expression than on production of IFN-γ or IL-21.

The most compelling result here is that skewing the hybrid-lineage cells towards a more committed Th1 phenotype in STAT3 TKO dramatically sped up clearance of parasitemia on reinfection. Supporting this interpretation, T-bet deficient mice, with less Th1-bias and more IFN-γ^−^IL-21^+^ T cells, were significantly slower at clearing a second infection. The mechanism of evolutionary pressure regulating this balance becomes clear in that the shift towards Th1 in STAT3 TKO animals, as well as the shift away from Th1 in the T-bet KO, both prolong high parasitemia and pathology in the first infection, as also shown for *P. berghei* ANKA. STAT3 has been previously suggested to be able to promote Tfh while inhibiting Th1 differentiation (Ray et al., 2014; Wu et al., 2015). However, the cytokine(s) responsible have not been identified (Findlay et al., 2010; Guthmiller et al., 2017; Gwyer Findlay et al., 2014; Hibbert et al., 2003; Kane et al., 2014; Ma et al., 2012b; Perez-Mazliah et al., 2015; Stumhofer et al., 2007).

Previous studies have demonstrated that the delicate balance of a *P. chabaudi* infected animal’s life or death is regulated by CD4 T cells, as is the case in other persistent infections such as tuberculosis and toxoplasmosis (Caruso et al., 1999; Denkers and Gazzinelli, 1998; Stephens et al., 2005). Animals deficient in either of the pro-Th1 factors IL-12 or IFN-γ or the regulatory cytokine IL-10, can all die of *P. chabaudi* infection even though it is a normally mild infection in mice (Li et al., 1999; Su and Stevenson, 2000, 2002). T-bet has been shown to be required for control of *P. berghei* ANKA parasitemia but is also essential for pathogenesis of experimental cerebral malaria (Oakley et al., 2013). While we did not see any mortality from an increase in Th1-type cells in the infected STAT3 TKO, mice that either had more (STAT3) or less (T-bet) IFN-γ^+^IL-21^−^ Th1-type cells had prolonged pathology. Therefore, while our data, and the human literature, suggest that a stronger Th1 response is beneficial for immunity to P. chabaudi, it remains to be tested if the combination of Th1/Tfh and regulatory cytokines into one cell type represents an evolutionary benefit, particularly in the first infection which is likely to drive evolution the most (next to pregnancy malaria). Further work considering the finely-tuned balance required to ensure host survival is needed to determine if this hybrid response is maladaptive.

In summary, persistent *Plasmodium* infection drives generation of a plastic mixed-lineage T cell with characteristics of both uncommitted Th1 (T-bet^int^) and pre-Tfh (CXCR5^int^), that is balanced by STAT3, Bcl6, Blimp-1 and T-bet, which coordinate the relative degree of antibody and IFN-γ responses for optimal pathogen control and host survival. While changing this balance towards Th1 in the first infection may prolong pathology, it promotes sterilizing immunity in the longer term, suggesting a potential direction for vaccine development.

## Acknowledgements

This work was supported by the NIH National Institute of Allergy and Infectious Diseases (R01AI08995304 (R.S., V.H.C.), R01AI135061 (R.S., V.H.C., F.A.); R01 AI132771 (A.L.D.); R01AI08995304S1 (R.S., V.H.C.) and F31AI126809 (V.H.C.) and the James W. McLaughlin and Jeane B. Kempner Fellowships (V.H.C., K.D.W.). We appreciate the expertise of Mark Griffin in the UTMB Microbiology & Immunology Flow Cytometry Core, and Gabriela M. Kaus and Margarita Ramirez for expert animal colony maintenance. We appreciate the scientific contribution of all Stephens and Joint Immunology Lab meeting members (Y. Cong, J. Sun, H. Hu, J.J. Endsley, R. Rajsbaum, L. Soong). Thanks to Roza I. Nurieva, John O’Shea, and Noah S. Butler for ideas, reagents and protocols.

## Author contributions

Conceptualization, V.H.C., R.S.; Methodology, V.H.C., R.S.; Investigation, V.H.C., K.D.W., F.A.; Writing-Original Draft, V.H.C.; Writing-Review & Editing, V.H.C., R.S., A.L.D.; Funding Acquisition, R.S., V.H.C., A.L.D.; Resources, A.L.D., R.S.

## STAR★Methods

### KEY RESOURCE TABLE

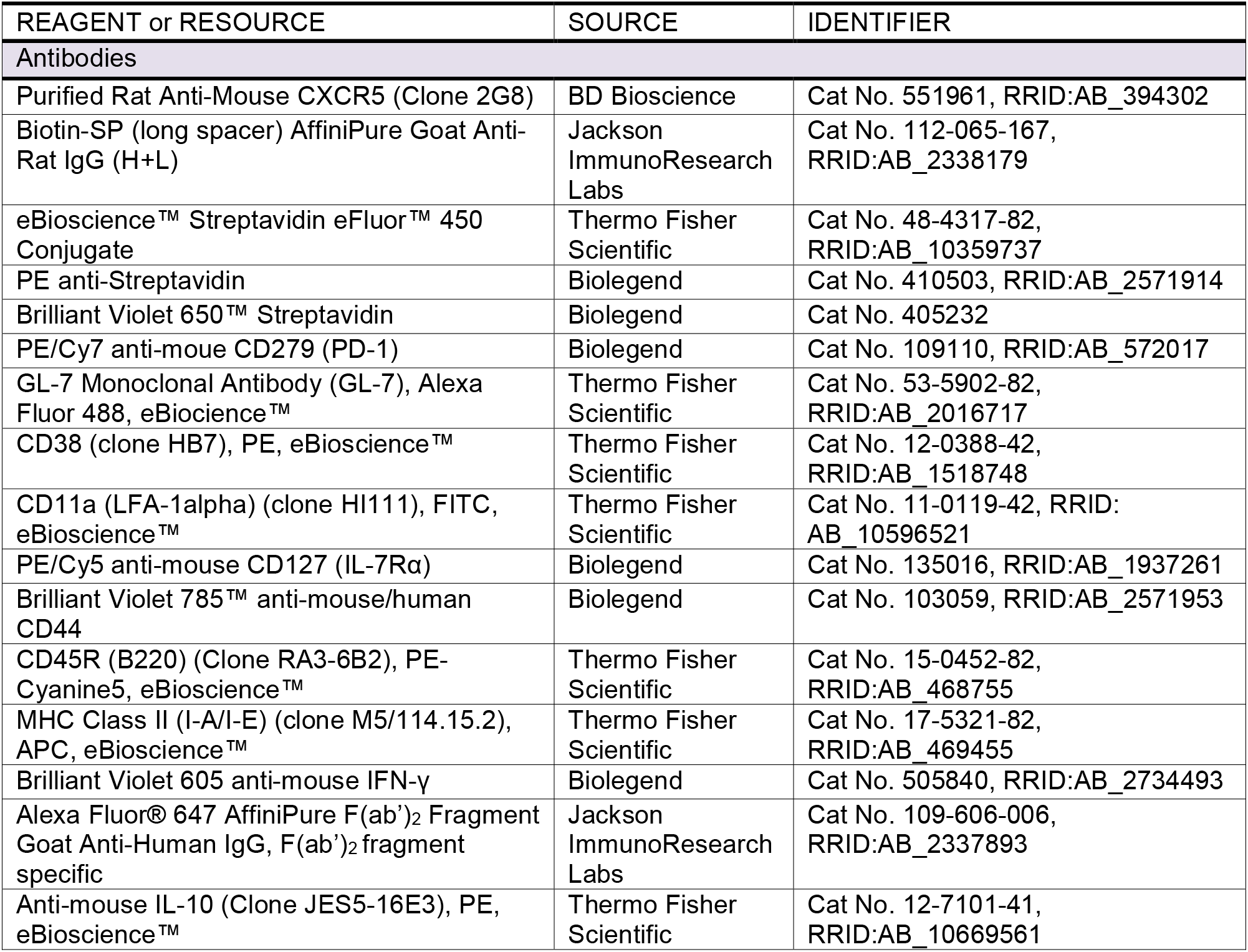
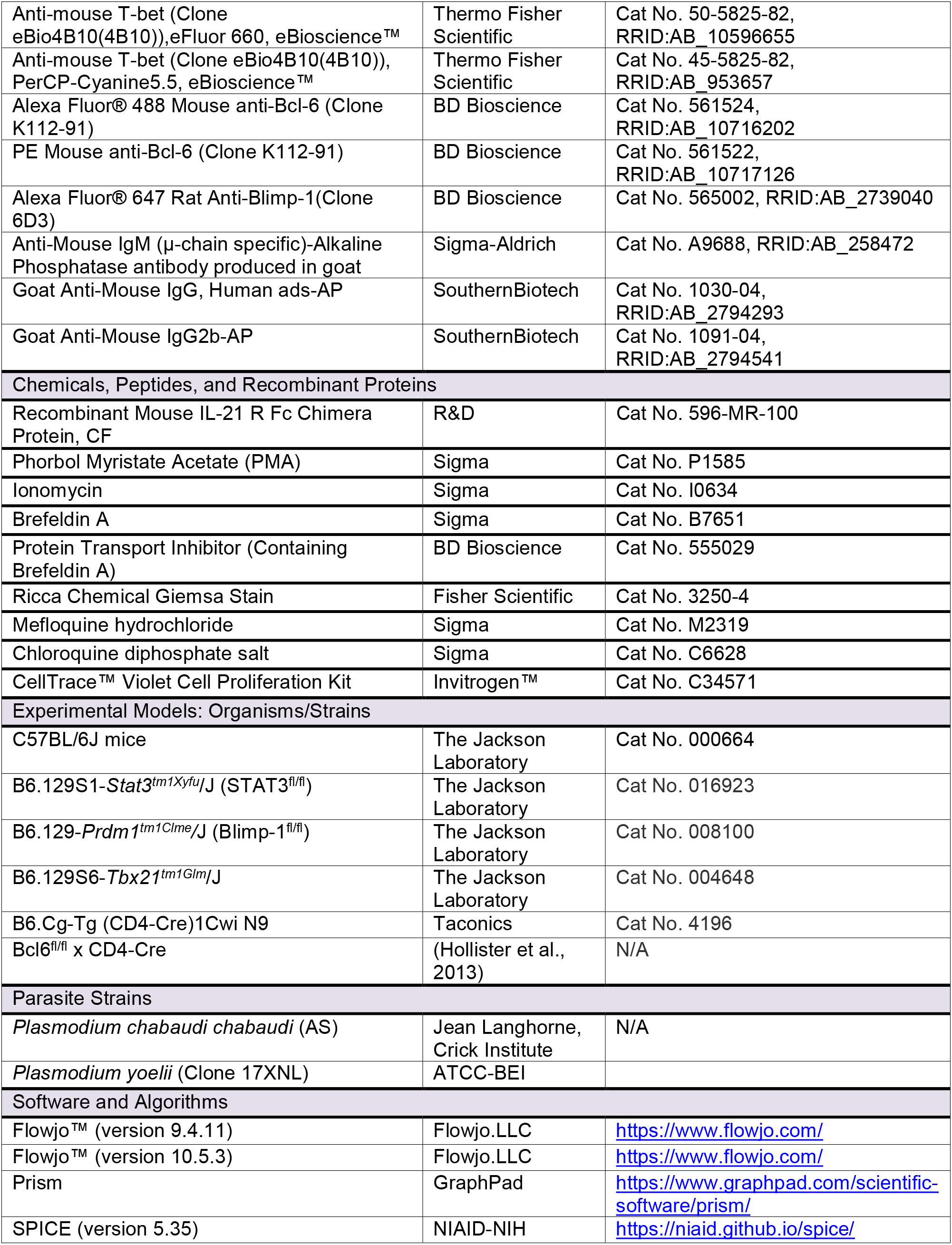

### CONTACT FOR REAGENT AND RESOURCE SHARING

Further information and requests for resources should be directed to, and will be fulfilled by the Lead Contact, Robin Stephens

### EXPERIMENTAL MODEL AND SUBJECT DETAILS

C57BL/6J (B6), B6.129S1-*Stat3*^*tm1Xyfu*^/J (*Stat3*^fl/fl^), B6.129-*Prdm1^tm1Clme^*/J (Blimp-1^fl/fl^), and B6.129S6-*Tbx21*^*tm1Glm*^/J (T-bet KO) mice were purchased from The Jackson Laboratory (Bar Harbor, ME) and bred to B6.Cg-Tg (CD4-Cre)1Cwi N9 mice from Taconic (Hudson, NY). *Bcl6*^fl/fl^ x CD4-Cre mice (Indiana University School of Medicine, Indianapolis, IN) were bred at UTMB. Six to twelve-week-old animals of both sexes were used for all experiments. All mice were maintained in our specific pathogen free animal facility with ad libitum access to food and water. All animal experiments were carried out in compliance with the protocol specifically approved for this study by the University of Texas Medical Branch Institutional Animal Care and Use Committee. Mice were infected i.p. with 10^5^ (or 10^7^ for re-infection) *Plasmodium chabaudi chabaudi* (AS; courtesy of Jean Langhorne (Francis Crick Institute, London, UK)) or 10^5^ *Plasmodium yoelii* (clone 17XNL; MR4/ATCC) infected red blood cells (iRBCs). Parasites were counted in thin blood smears stained with Giemsa (Sigma, St. Louis, MO) by light microscopy. In some experiments, mice were treated with mefloquine hydrochloride (MQ, 4mg/kg body weight, Sigma, St. Louis, MO) by oral gavage daily five times or until the mice were euthanized. In some experiments (STAT3 TKO) mice were treated with 50 mg/kg body weight per animal of Chloroquine (CQ) in Saline (both from Sigma) every other day for a total of three times, starting 10 weeks p.i..

## METHOD DETAILS

### Flow Cytometry and adoptive transfer

Single-cell suspensions from spleens were made in Hank’s Balanced Salt Solution (Gibco, Lifetechnologies, Grand Island, NY), with added HEPES (Sigma), followed by red blood cell lysis buffer (eBioscience, San Diego, CA). Multicolor panels including CXCR5 were stained in PBS + 0.5% BSA + 0.1% sodium azide + 2% Normal Mouse Serum (NMS) and 2% FBS (Sigma, St. Louis, MO). Rat anti-mouse purified CXCR5 (2G8, BDbioscience, San Jose, CA, 1 hr, 4°C) was followed by biotin-conjugated AffiniPure Goat anti-rat (H+L, Jackson Immunoresearch, West Grove, PA, 30 min, 4°C) followed by Streptavidin-eFluor 450, –PE or – Brilliant Violet 650 (BV650). As described in Crotty et al, the third step included the other antibodies-combinations of FITC–, phycoerythrin (PE)–, Peridinin Chlorophyll Protein Complex (PerCP)-Cyanine (Cy)5.5, PE/ Cyanine 7 (Cy7), Allophycocyanin (APC) monoclonal antibodies (all from eBioscience, San Diego, CA), and CD127-PE/Cy5, CD44-Brilliant Violet 785 (Biolegend, San Diego, CA). For B cell staining we used B220-PE/Cy5, MHC-II(I-A/I-E)-APC, CD38-PE, GL-7-FITC (all from eBioscience, San Diego, CA). For intracellular staining, total cells were stimulated for 2 h with phorbol myristate acetate (PMA, 50 ng/mL), Ionomycin (500 ng/mL), and Brefeldin A (10 μg/mL, all from Sigma) in complete Iscove’s Media (cIMDM+, +10% FBS, +2mM L-glutamine, +0.5 mM sodium pyruvate, +100 U/ml penicillin, +100ug/ml streptomycin, +50 μM 2-β-Mercaptoethanol ((ME); all from Gibco, Lifetechnologies). Cells were fixed in 2% paraformaldehyde (Sigma), permeabilized using Permeabilization buffer (10X Permeabilization buffer, eBioscience) and incubated for 40 minutes with anti-IFN-γ-Brilliant Violet 605 (XMG1.2), T-bet-efluor 660 or -PerCP-Cy5.5 (eBio4B10, eBioscience), Bcl6-Alexa Fluor 488 or –PE (K112-91), and/or Blimp-1-Alexa Fluor 647 (6D3, BDbioscience). For IL-21 staining, cells were incubated with recombinant mouse IL-21R-Fc chimera (1 μg, 40 min., R&D systems, Minneapolis, MN) and washed twice in Perm buffer, followed by Alexa Fluor 647 goat anti-human IgG F(ab’)₂ (0.3 μg, 30 min, Jackson ImmunoResearch, West Grove, PA) in Perm buffer. After three washes in FACS buffer, cells were collected on a LSRII Fortessa at the UTMB Flow Cytometry and analyzed in FlowJo versions 9.4.11, 10.5.3 (TreeStar, Ashland, OR). Compensation was performed in FlowJo using single CD4 stained splenocytes.

Cell Trace Violet (CTV, Invitrogen) staining of splenocytes was done in calcium- and magnesium-free PBS at 10^7^ cells/ml with 5μM CTV for 10 minutes at 37°C with shaking, then quenched with Fetal Calf Serum. After washing, 2 x10^6^ cells were transferred into each mouse i.p. In house Brefeldin A solution was used in all figures, except Figures 4B and S7A STAT3 TKO in which we used commercial Brefeldin (GolgiPlug).

### ELISA

Serum samples were obtained on the indicated days by bleeding mice from the tail vein under a heat lamp. Nunc-Immuno Plates (MaxiSorp™) were coated with whole freeze-thaw parasite lysate (transfer from N_2_(l) to 37°C, 4-5 times (Guthmiller et al., 2017). Plates were blocked with 2.5% BSA + 5%FCS in PBS. Bound antibody was detected using Alkaline Phosphatase (AP)-conjugated goat anti-mouse IgM (Sigma), IgG and IgG2b (Southern Biotech, Brimingham, AL) which was revealed with a 4-Nitrophenyl phosphate disodium salt hexahydrate (PNPP, Sigma) solution (1 mg/ml). Plates were analyzed with a FLUOstar Omega plate reader (BMG Labtech, Cary, NC).

### Statistics

Statistical analysis was performed in Prism (GraphPad, La Jolla, CA) using Student’s *t*-test. *p* < 0.05 was accepted as a statistically significant difference, * *p* ≤0.05, \*\**p* ≤0.01, \*\*\**p* ≤0.001, \*\*\*\**p* ≤0.001. Boolean gating analysis and Pie graphs were performed in SPICE software version 5.35 (http://exon.niaid.nih.gov/spice/).

---

**Figure S1.**
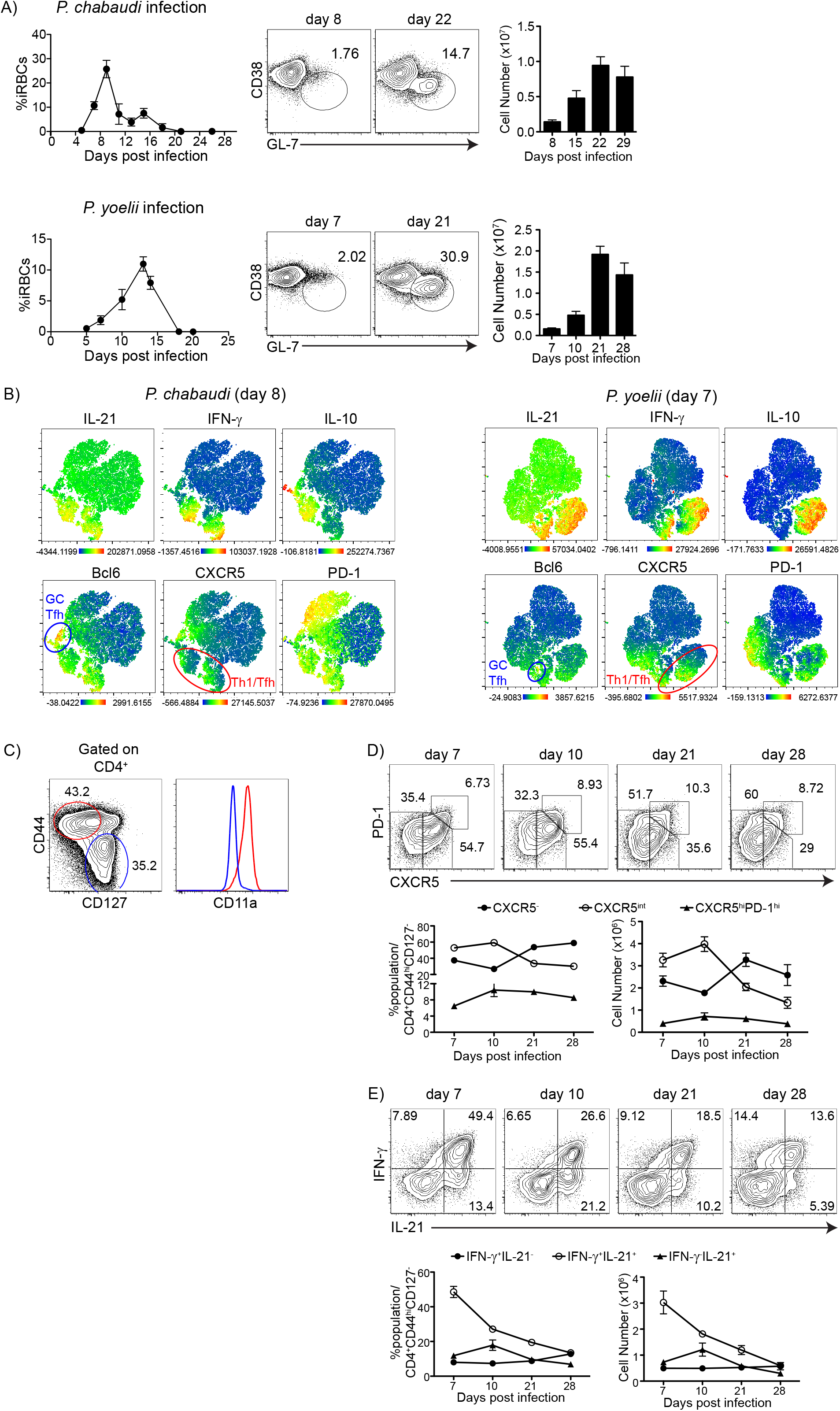
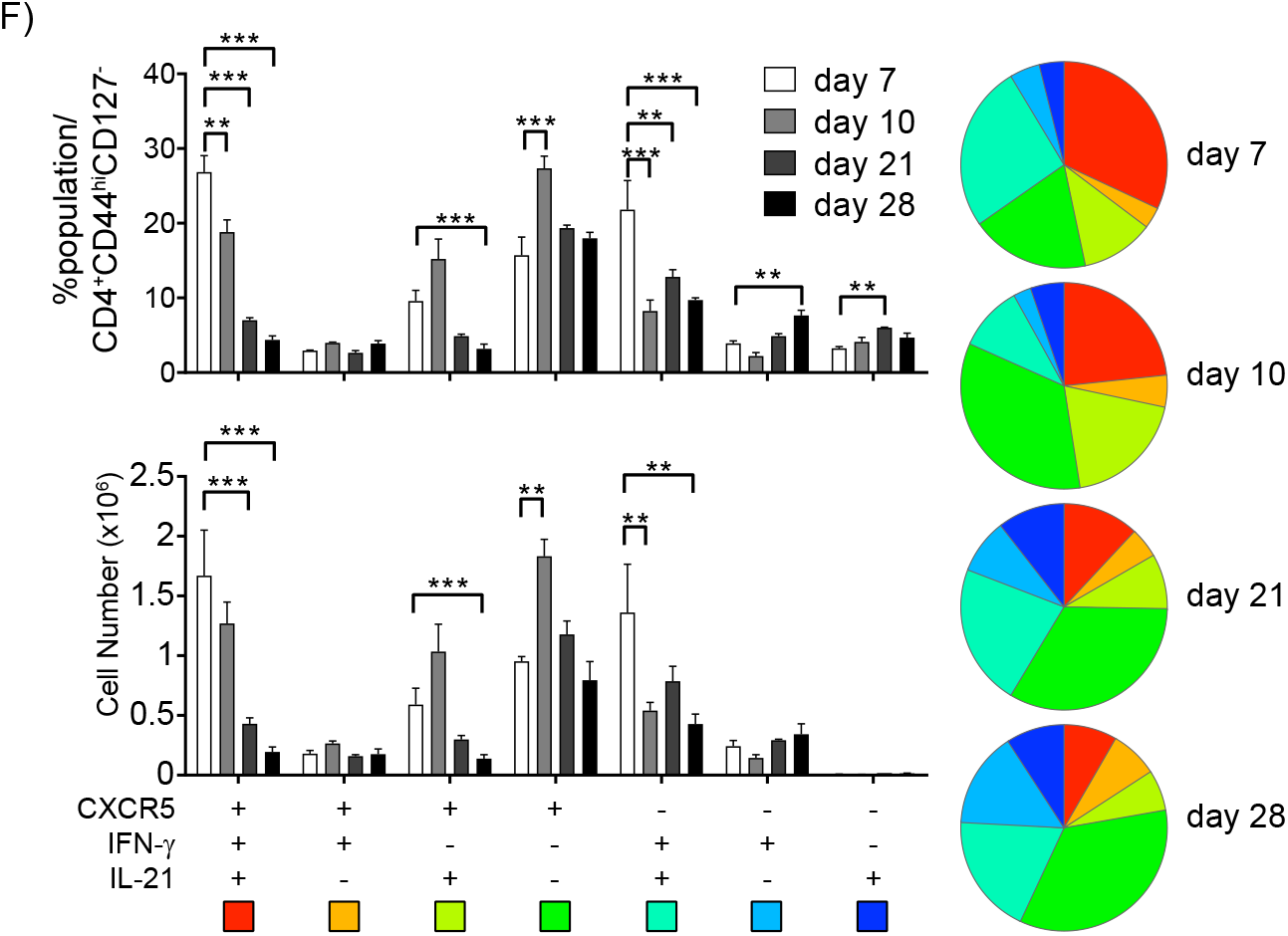
T helper differentiation during *P. chabaudi* and *P. yoelii* 17XNL infection. C57BL/6J mice were infected and splenocytes were analyzed on days indicated. **(A)** Parasitemia curves, and contour plots showing expression of CD38 and GL-7 in B cells (B220^hi^MHC-II^hi^). Bar graph shows numbers of GC B cells at indicated days. **(B)** Density t-SNE plots of CD4^+^ T cells from C57BL/6J mice infected with *P. chabaudi* at day 8 p.i. or *P. yoelii* at day 7 p.i. Plots show 10^5^ representative T cells from each of 3 mice, concatenated and overlaid with the expression of selected markers. **(C)** Contour plots show CD44, CD127, and CD11a expression in CD4^+^ T cells at day 8 of *P. chabaudi* infection. Majority of CD44^hi^CD127^−^CD4^+^ T cells are CD11a^hi^. Contour plots show expression of **(D)** PD-1 and CXCR5 and **(E)** IFN-γ and IL-21 in CD4 Teff during *P. yoelii* infection. Right line graphs show percentage (left) and numbers (right). **(F)** Boolean gating of CXCR5, IFN-γ, and IL-21 expression of CD4 Teff in *P. yoelii* infection. Pie charts show the distribution of all the subsets on each day. Bar graphs show the percentages and cell numbers of the subsets on each day. Data representative of 2 experiments with 3 mice/group.

**Figure S2.**
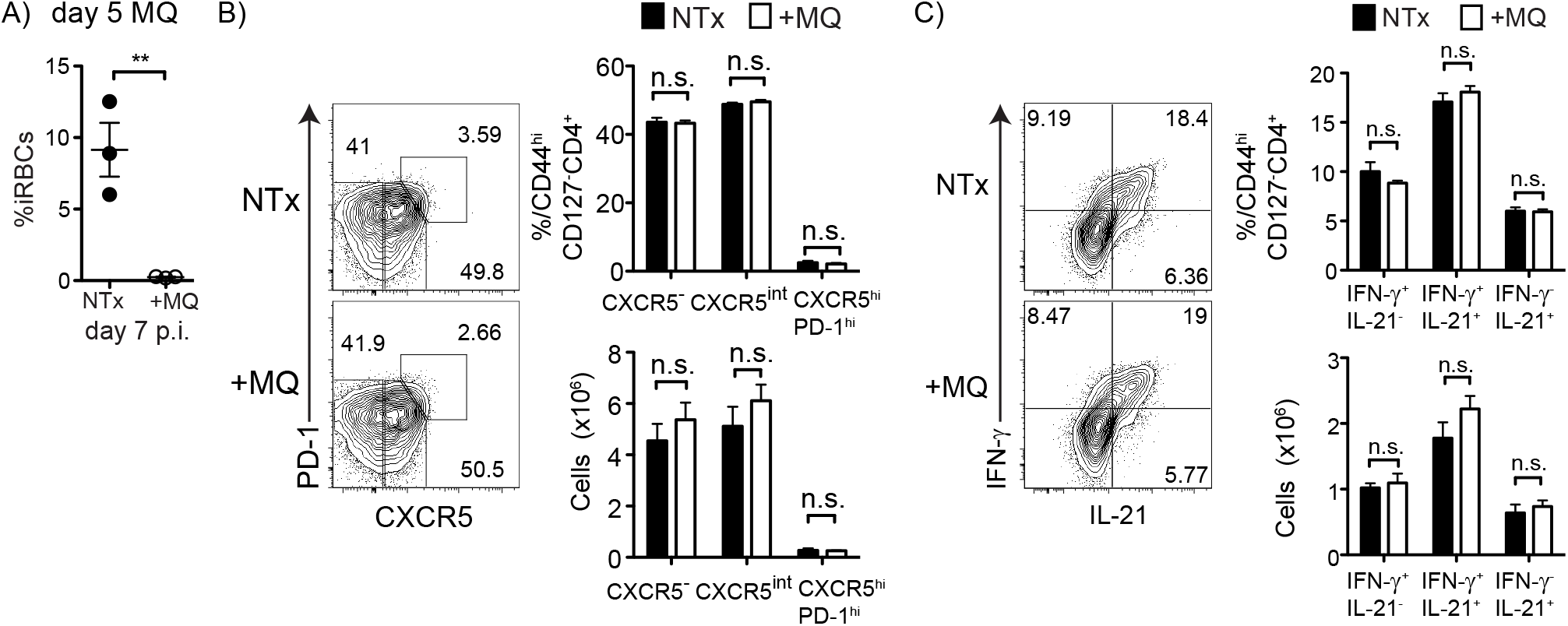
Day 5 mefloquine treatment has no effect on hybrid Th1/Tfh cell differentiation. C57BL/6J mice were infected, and one group was treated with mefloquine starting day 5 and splenocytes were analyzed at day 7 p.i. (**A**) Parasitemia on day 7 p.i. from untreated (NTx, black filled circles) and treated (+MQ, open circles) groups. Contour plots show expression of **B**) PD-1 and CXCR5, and C) IFN-γ and IL-21 in Teff. Bar graphs show fraction of Teff and numbers of subsets. Data representative of 2 experiments with 3 mice/group.

**Figure S3.**
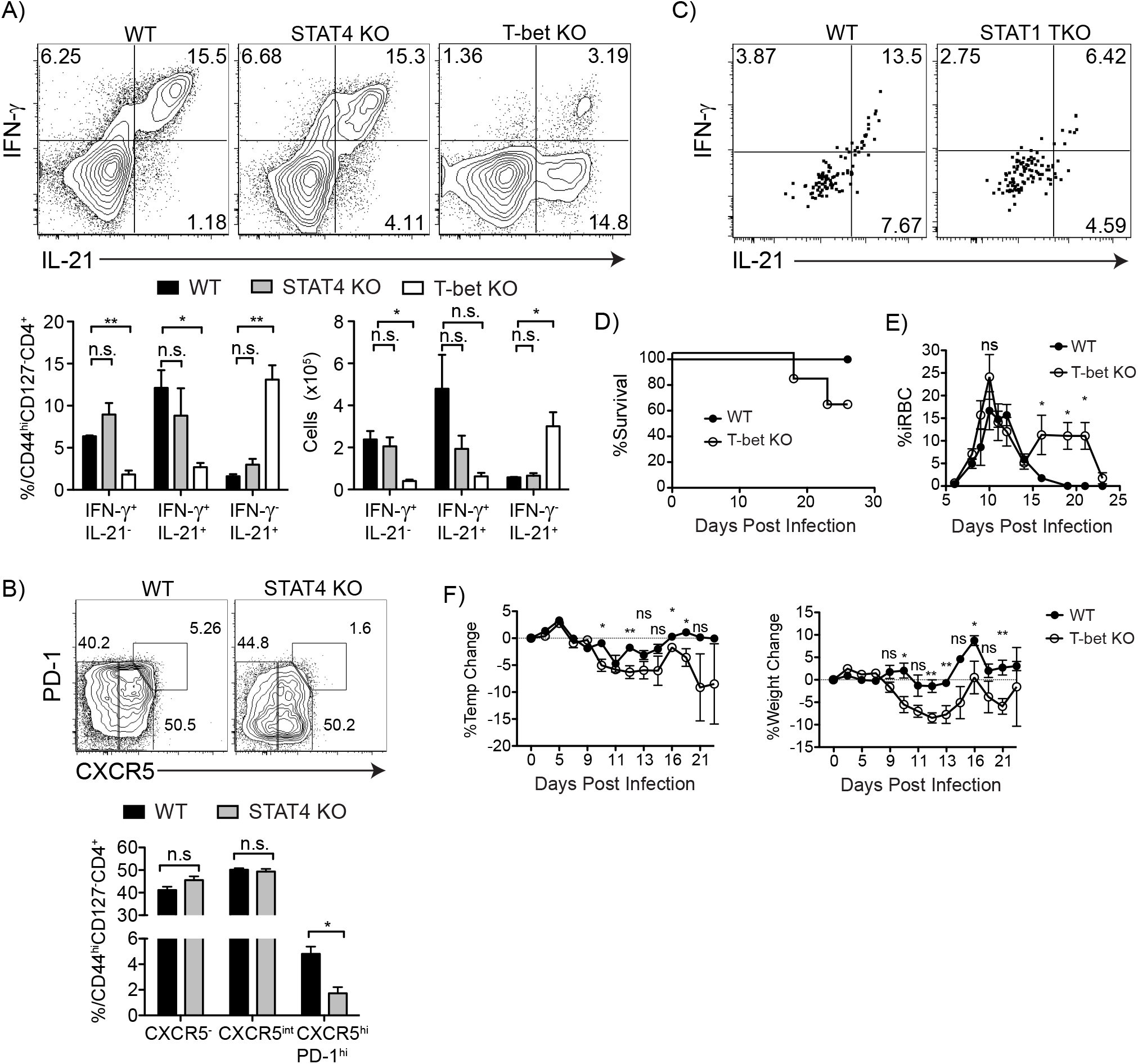
T-bet, but not STAT4 nor STAT1, is required for IFN-γ production by hybrid Th1/Tfh. (**A**) C57BL/6J (WT), STAT4 KO, and T-bet KO mice were infected and splenocytes were analyzed at day 7 p.i. Contour plots and bar graphs show expression of IFN-γ and IL-21 in Teff from WT (black bar), STAT4 KO (gray bar) and T-bet KO (white bar). **(B)** Contour plots show PD-1 and CXCR5 expression in Teff from WT and STAT4 KO mice at day 7 p.i. Below, bar graphs show percentages. **(C)** Splenocytes from uninfected *Stat1*^fl/fl^CD4^Cre^ (STAT1 TKO) or *Stat1*^fl/fl^ (WT) were labeled with cell trace violet (CTV) and adoptively transferred into Ly5.1 (CD45.1) congenic mice, followed by *P. chabaudi* infection. Contour plots show expression of IFN-γ and IL-21 in CTV^−^ gated Teff on day 8 p.i. **(D)** Survival curve and **(E)** Parasitemia of WT (filled circles) and T-bet KO (open circles). **(F)** Temperature and weight loss of infected WT (filled circles) and T-bet KO (open circles). Data representative of 2 experiments with 3-8 mice/group for (**A, D, E, and F**) and 1 experiment with 4-5 mice/group for (C).

**Figure S4.**
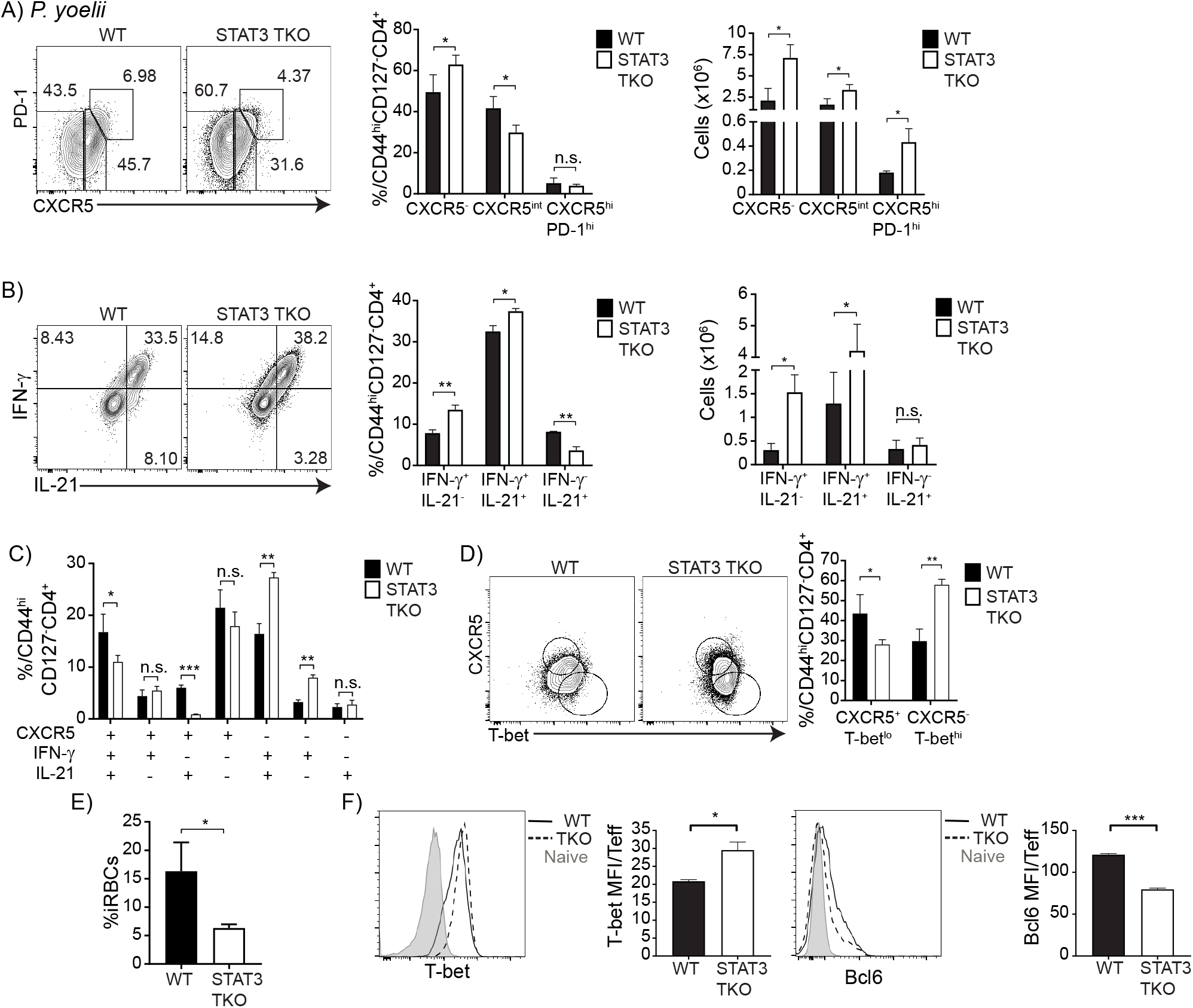
*P. yoelii*-infected STAT3 TKO mice show similar T cell differentiation than *P. chabaudi* infection. *Stat3*^fl/fl^CD4^Cre^ (TKO) and *Stat3*^fl/fl^ (WT) animals were infected and splenocytes were analyzed at day 10 p.i. Contour plots show fraction of subsets using **(A)** PD-1 and CXCR5, and **(B)** IFN-γ and IL-21 gated on Teff. Bar graphs show percentages and numbers of subsets. **(C)** Boolean gating of CXCR5, IFN-γ, and IL-21 within WT (black bars) and STAT3 TKO (white bars) Teff. **(D)** Contour plots show expression of CXCR5 and T-bet in Teff. Bar graph shows percentages CXCR5^+^T-bet^lo^ and CXCR5^−^T-bet^hi^ subsets. **(E)** Parasitemia of WT (black filled dots) and STAT3 TKO (open circles) animals on day 10 p.i. **(F)** Histograms showing T-bet (left) and Bcl6 (right) expression in Teff from STAT3 TKO (dotted line) and WT (black line) animals, and naive (gray filled line) cells. Bar graphs shows average MFI of T-bet and Bcl6. Data representative of 1 experiment with 2-3 mice/group.

**Figure S5.**
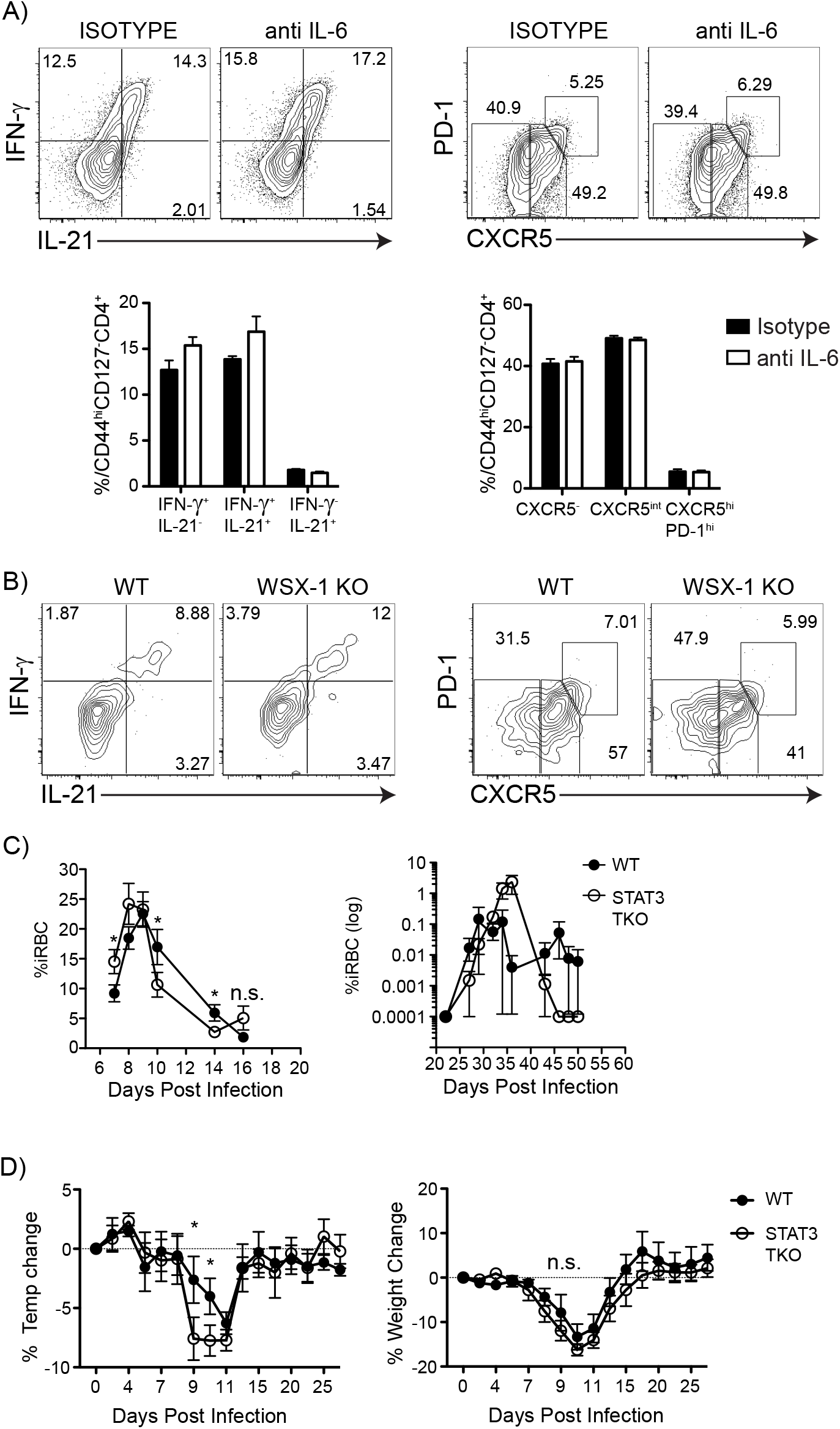
IL-6 blockade nor WSX-1 deficiency do not recapitulate STAT3 TKO mice phenotype. **(A)** C57BL/6J mice (n=5/group) were infected and either treated with anti-IL-6 or isotype antibodies. Contour plots show expression of IFN-γ and IL-21, and PD-1 and CXCR5 in Teff at day 7 p.i. **(B)** Splenocytes from uninfected WSX-1 KO or C57BL/6J were labeled with cell trace violet (CTV) and adoptively transferred into Ly5.1 (CD45.1) congenic mice, followed by *P. chabaudi* infection. Contour plots show expression of IFN-γ/IL-21 and PD-1/CXCR5 in CTV^−^ gated Teff on day 8 p.i. *STAT3*^fl/fl^CD4^Cre^ (TKO) and *STAT3*^fl/fl^ (WT) animals were infected and followed for 30 days. **(C)** Parasitemia of WT (black filled dots) and STAT3 TKO (open circles) animals. **(D)** Temperature and weight loss of WT (black filled dots) and STAT3 TKO (open circles) animals are shown in percentages. (**A, B**) Data representative of 1 experiment, 3-5 mice/group. (**C, D**) Data representative of 3 experiments, 3-8 mice/group.

**Figure S6.**
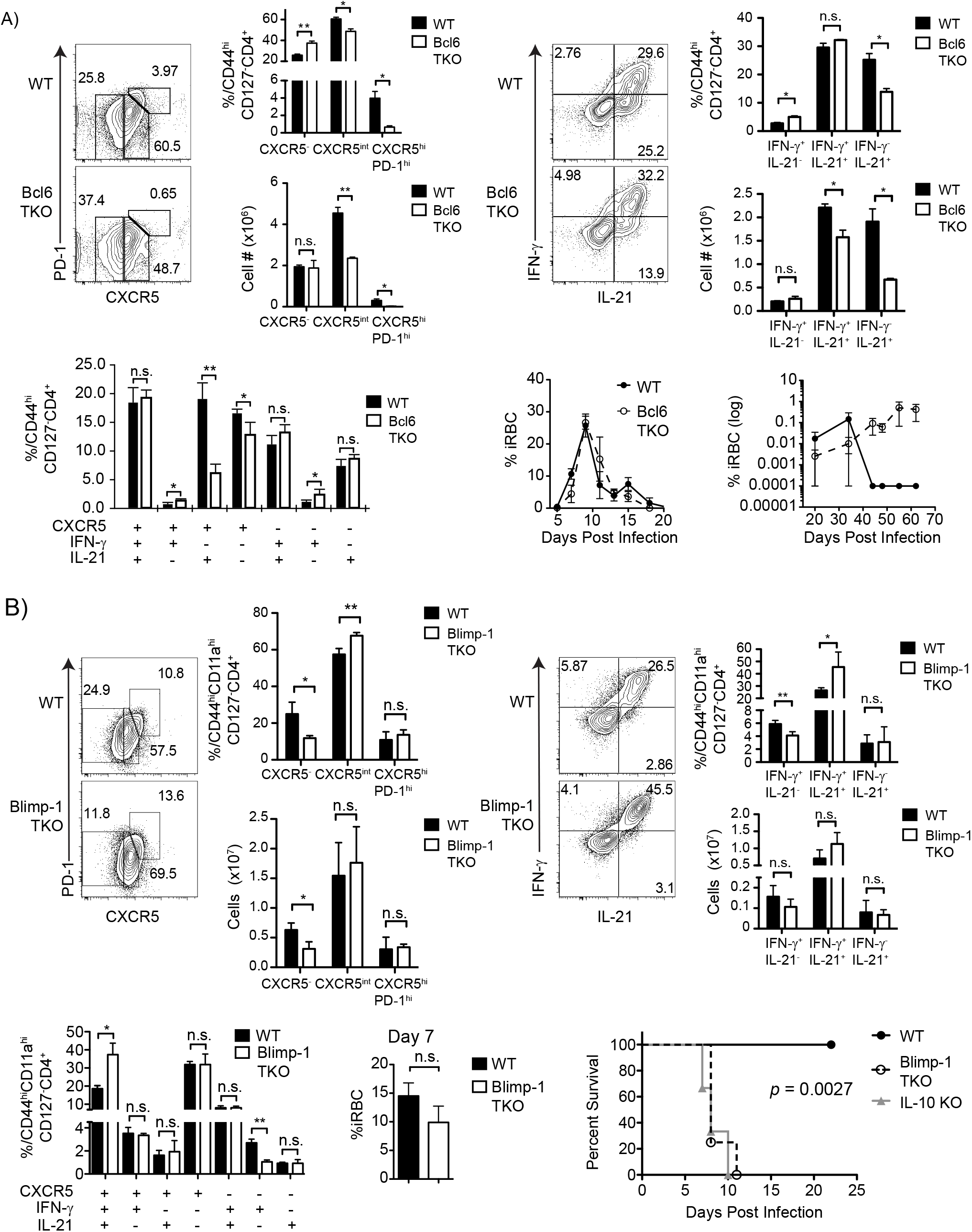
Roles of Bcl6 and Blimp in T cell differentiation during *P. chabaudi* infection. **(A)** *Bcl6*^fl/fl^CD4^Cre^ (TKO) and *Bcl6*^fl/fl^ (WT) animals were infected and splenocytes were analyzed at day 7 p.i. Contour plots and bar graphs show expression of subsets of PD-1/CXCR5, and IFN-γ/IL-21 in Teff, GC Tfh are CXCR5^hi^PD-1^hi^. Boolean analysis of CXCR5, IFN-γ and IL-21 within WT (black) and Bcl6 TKO (white) Teff. Parasitemia of WT (black filled circles) and Bcl6 TKO (open circles) animals. **(B)** *Blimp-1*^fl/fl^CD4^Cre^ (TKO) and *Blimp-1*^fl/fl^ (WT) animals were infected and splenocytes were analyzed at day 7 p.i. Contour plots and bar graphs show subsets of PD-1/CXCR5, and IFN-γ/IL-21 gated on Teff. Boolean analysis of CXCR5, IFN-γ and IL-21 within WT (black bars) and Blimp-1 TKO (white bars) Teff. Parasitemia of WT (black bars) and Blimp-1 TKO (white bars) animals at day 7 p.i. Survival of WT (black circles), Blimp-1 TKO (open circles) and IL-10 KO (filled gray triangles). Data representative of 3 experiments, 3-4 mice/group.

**Figure S7.**
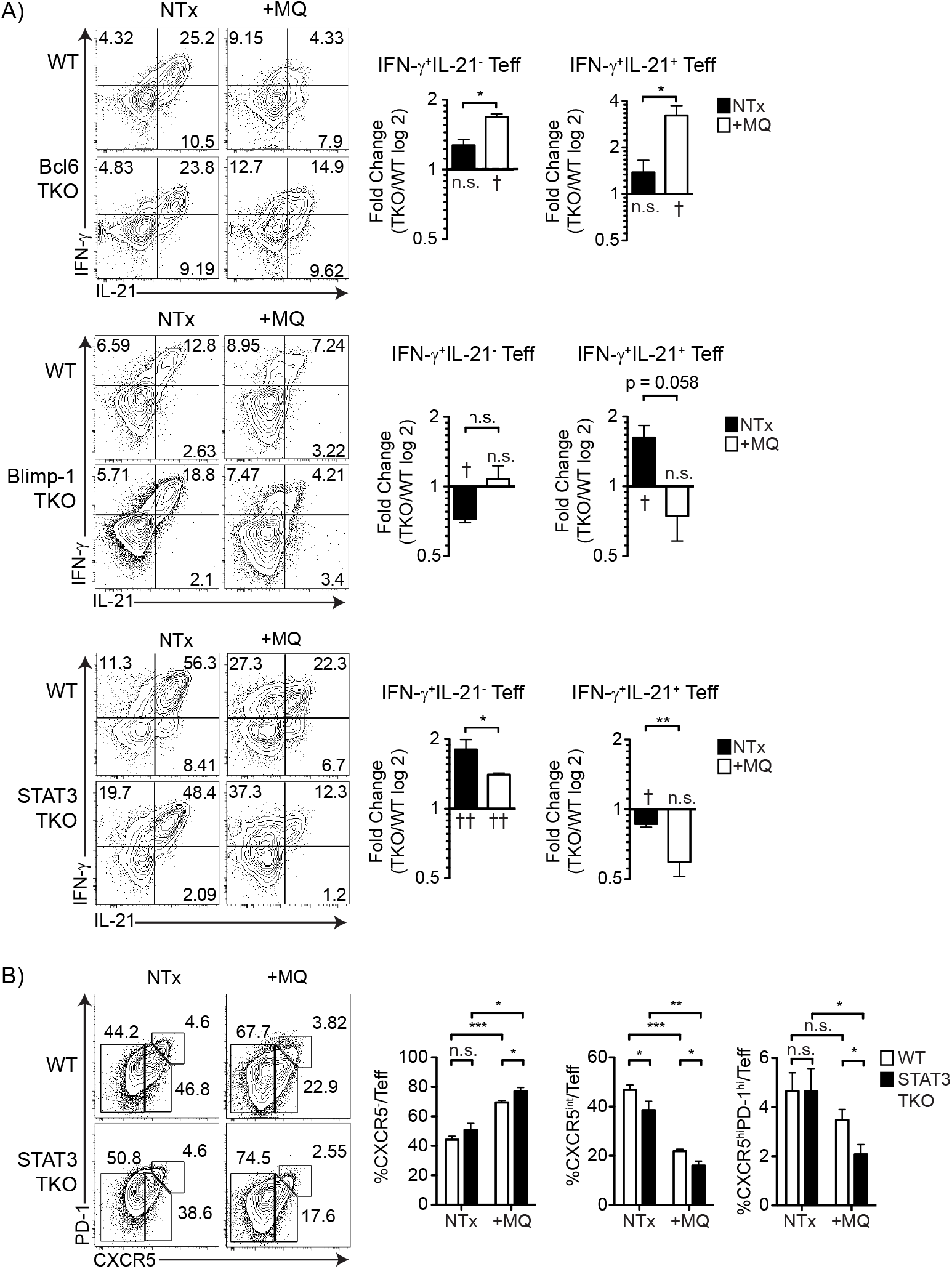
Response to *P. chabaudi* infection in drug-cured Bcl6, Blimp-1, and STAT3 TKO mice. TKO and WT animals were infected. Animals were given mefloquine (+MQ) starting on day 3 p.i. or left untreated (NTx). Splenocytes were harvested and analyzed by flow cytometry at day 7 p.i. **(A)** Contour plots show expression of IFN-γ and IL-21 in Teff. Bar graphs show average of fold change (%TKO/%WT) in a log2 scale of percentages of IFN-γ^+^IL-21^−^ and IFN-γ^+^IL-21^−^ Teff from NTx (black bars) and +MQ (white bars) from Bcl6 TKO (top), Blimp-1 TKO (middle), and STAT3 TKO (bottom). Intracellular cytokine staining in STAT3 TKO was the only one done with commercially prepared secretion inhibitor. **(B)** Contour plots show expression of PD-1 and CXCR5 in Teff from STAT3 TKO and WT mice. Bar graphs show percentages. Data representative of 2 experiments with 3 mice/group. † *p* < 0.05, †† *p* < 0.01 are statistical comparison of WT and TKO mice within NTx and +MQ groups.

## Reference

Hollister, K., Kusam, S., Wu, H., Clegg, N., Mondal, A., Sawant, D.V., and Dent, A.L. (2013). Insights into the role of Bcl6 in follicular Th cells using a new conditional mutant mouse model. J Immunol 191, 3705–3711.

## References

Afkarian, M., Sedy, J.R., Yang, J., Jacobson, N.G., Cereb, N., Yang, S.Y., Murphy, T.L., and Murphy, K.M. (2002). T-bet is a STAT1-induced regulator of IL-12R expression in naive CD4+ T cells. Nat Immunol 3, 549–557.

Amante, F.H., and Good, M.F. (1997). Prolonged Th1-like response generated by a Plasmodium yoelii-specific T cell clone allows complete clearance of infection in reconstituted mice. Parasite Immunol 19, 111–126.

Batten, M., Ramamoorthi, N., Kljavin, N.M., Ma, C.S., Cox, J.H., Dengler, H.S., Danilenko, D.M., Caplazi, P., Wong, M., Fulcher, D.A., et al. (2010). IL-27 supports germinal center function by enhancing IL-21 production and the function of T follicular helper cells. J Exp Med 207, 2895–2906.

Bentebibel, S.E., Lopez, S., Obermoser, G., Schmitt, N., Mueller, C., Harrod, C., Flano, E., Mejias, A., Albrecht, R.A., Blankenship, D., et al. (2013). Induction of ICOS+CXCR3+CXCR5+ TH cells correlates with antibody responses to influenza vaccination. Sci Transl Med 5, 176ra132.

Carpio, V.H., Opata, M.M., Montanez, M.E., Banerjee, P.P., Dent, A.L., and Stephens, R. (2015). IFN-gamma and IL-21 Double Producing T Cells Are Bcl6-Independent and Survive into the Memory Phase in Plasmodium chabaudi Infection. PLoS One 10, e0144654.

Caruso, A.M., Serbina, N., Klein, E., Triebold, K., Bloom, B.R., and Flynn, J.L. (1999). Mice deficient in CD4 T cells have only transiently diminished levels of IFN-gamma, yet succumb to tuberculosis. J Immunol 162, 5407–5416.

Choi, Y.S., Eto, D., Yang, J.A., Lao, C., and Crotty, S. (2013). Cutting edge: STAT1 is required for IL-6-mediated Bcl6 induction for early follicular helper cell differentiation. J Immunol 190, 3049–3053.

Cimmino, L., Martins, G.A., Liao, J., Magnusdottir, E., Grunig, G., Perez, R.K., and Calame, K.L. (2008). Blimp-1 attenuates Th1 differentiation by repression of ifng, tbx21, and bcl6 gene expression. J Immunol 181, 2338–2347.

Crawford, A., Angelosanto, J.M., Kao, C., Doering, T.A., Odorizzi, P.M., Barnett, B.E., and Wherry, E.J. (2014). Molecular and transcriptional basis of CD4(+) T cell dysfunction during chronic infection. Immunity 40, 289–302.

Crotty, S. (2014). T follicular helper cell differentiation, function, and roles in disease. Immunity 41, 529–542.

Curtis, M.M., Rowell, E., Shafiani, S., Negash, A., Urdahl, K.B., Wilson, C.B., and Way, S.S. (2010). Fidelity of pathogen-specific CD4+ T cells to the Th1 lineage is controlled by exogenous cytokines, interferon-gamma expression, and pathogen lifestyle. Cell Host Microbe 8, 163–173.

Denkers, E.Y., and Gazzinelli, R.T. (1998). Regulation and function of T-cell-mediated immunity during Toxoplasma gondii infection. Clin Microbiol Rev 11, 569–588.

Elsaesser, H., Sauer, K., and Brooks, D.G. (2009). IL-21 is required to control chronic viral infection. Science 324, 1569–1572.

Fahey, L.M., Wilson, E.B., Elsaesser, H., Fistonich, C.D., McGavern, D.B., and Brooks, D.G. (2011). Viral persistence redirects CD4 T cell differentiation toward T follicular helper cells. J Exp Med 208, 987–999.

Fang, D., Cui, K., Mao, K., Hu, G., Li, R., Zheng, M., Riteau, N., Reiner, S.L., Sher, A., Zhao, K., et al. (2018). Transient T-bet expression functionally specifies a distinct T follicular helper subset. J Exp Med 215, 2705–2714.

Findlay, E.G., Greig, R., Stumhofer, J.S., Hafalla, J.C., de Souza, J.B., Saris, C.J., Hunter, C.A., Riley, E.M., and Couper, K.N. (2010). Essential role for IL-27 receptor signaling in prevention of Th1-mediated immunopathology during malaria infection. J Immunol 185, 2482–2492.

Freitas do Rosario, A.P., Lamb, T., Spence, P., Stephens, R., Lang, A., Roers, A., Muller, W., O’Garra, A., and Langhorne, J. (2012). IL-27 promotes IL-10 production by effector Th1 CD4+ T cells: a critical mechanism for protection from severe immunopathology during malaria infection. J Immunol 188, 1178–1190.

Guthmiller, J.J., Graham, A.C., Zander, R.A., Pope, R.L., and Butler, N.S. (2017). Cutting Edge: IL-10 Is Essential for the Generation of Germinal Center B Cell Responses and Anti-Plasmodium Humoral Immunity. J Immunol 198, 617–622.

Gwyer Findlay, E., Villegas-Mendez, A., O’Regan, N., de Souza, J.B., Grady, L.M., Saris, C.J., Riley, E.M., and Couper, K.N. (2014). IL-27 receptor signaling regulates memory CD4+ T cell populations and suppresses rapid inflammatory responses during secondary malaria infection. Infect Immun 82, 10–20.

Hale, J.S., Youngblood, B., Latner, D.R., Mohammed, A.U., Ye, L., Akondy, R.S., Wu, T., Iyer, S.S., and Ahmed, R. (2013). Distinct memory CD4+ T cells with commitment to T follicular helper- and T helper 1-cell lineages are generated after acute viral infection. Immunity 38, 805–817.

Hansen, D.S., Obeng-Adjei, N., Ly, A., Ioannidis, L.J., and Crompton, P.D. (2017). Emerging concepts in T follicular helper cell responses to malaria. Int J Parasitol 47, 105–110.

Hibbert, L., Pflanz, S., De Waal Malefyt, R., and Kastelein, R.A. (2003). IL-27 and IFN-alpha signal via Stat1 and Stat3 and induce T-Bet and IL-12Rbeta2 in naive T cells. J Interferon Cytokine Res 23, 513–522.

Hsieh, C.S., Macatonia, S.E., Tripp, C.S., Wolf, S.F., O’Garra, A., and Murphy, K.M. (1993). Development of TH1 CD4+ T cells through IL-12 produced by Listeria-induced macrophages. Science 260, 547–549.

Hunt, P., Cravo, P.V., Donleavy, P., Carlton, J.M., and Walliker, D. (2004). Chloroquine resistance in Plasmodium chabaudi: are chloroquine-resistance transporter (crt) and multi-drug resistance (mdr1) orthologues involved? Mol Biochem Parasitol 133, 27–35.

Johnston, R.J., Poholek, A.C., DiToro, D., Yusuf, I., Eto, D., Barnett, B., Dent, A.L., Craft, J., and Crotty, S. (2009). Bcl6 and Blimp-1 are reciprocal and antagonistic regulators of T follicular helper cell differentiation. Science 325, 1006–1010.

Kane, A., Deenick, E.K., Ma, C.S., Cook, M.C., Uzel, G., and Tangye, S.G. (2014). STAT3 is a central regulator of lymphocyte differentiation and function. Curr Opin Immunol 28, 49–57.

Khader, S.A., Bell, G.K., Pearl, J.E., Fountain, J.J., Rangel-Moreno, J., Cilley, G.E., Shen, F., Eaton, S.M., Gaffen, S.L., Swain, S.L., et al. (2007). IL-23 and IL-17 in the establishment of protective pulmonary CD4+ T cell responses after vaccination and during Mycobacterium tuberculosis challenge. Nat Immunol 8, 369–377.

Li, C., Corraliza, I., and Langhorne, J. (1999). A defect in interleukin-10 leads to enhanced malarial disease in Plasmodium chabaudi chabaudi infection in mice. Infect Immun 67, 4435–4442.

Li, C., Sanni, L.A., Omer, F., Riley, E., and Langhorne, J. (2003). Pathology of Plasmodium chabaudi chabaudi infection and mortality in interleukin-10-deficient mice are ameliorated by anti-tumor necrosis factor alpha and exacerbated by anti-transforming growth factor beta antibodies. Infect Immun 71, 4850–4856.

Li, L., Jiang, Y., Lao, S., Yang, B., Yu, S., Zhang, Y., and Wu, C. (2016). Mycobacterium tuberculosis-Specific IL-21+IFN-gamma+CD4+ T Cells Are Regulated by IL-12. PLoS One 11, e0147356.

Lonnberg, T., Svensson, V., James, K.R., Fernandez-Ruiz, D., Sebina, I., Montandon, R., Soon, M.S., Fogg, L.G., Nair, A.S., Liligeto, U., et al. (2017). Single-cell RNA-seq and computational analysis using temporal mixture modelling resolves Th1/Tfh fate bifurcation in malaria. Sci Immunol 2.

Luty, A.J., Lell, B., Schmidt-Ott, R., Lehman, L.G., Luckner, D., Greve, B., Matousek, P., Herbich, K., Schmid, D., Migot-Nabias, F., et al. (1999). Interferon-gamma responses are associated with resistance to reinfection with Plasmodium falciparum in young African children. J Infect Dis 179, 980–988.

Ma, C.S., Avery, D.T., Chan, A., Batten, M., Bustamante, J., Boisson-Dupuis, S., Arkwright, P.D., Kreins, A.Y., Averbuch, D., Engelhard, D., et al. (2012a). Functional STAT3 deficiency compromises the generation of human T follicular helper cells. Blood 119, 3997–4008.

May, J., Lell, B., Luty, A.J., Meyer, C.G., and Kremsner, P.G. (2000). Plasma interleukin-10:Tumor necrosis factor (TNF)-alpha ratio is associated with TNF promoter variants and predicts malarial complications. J Infect Dis 182, 1570–1573.

McDermott, D.S., and Varga, S.M. (2011). Quantifying antigen-specific CD4 T cells during a viral infection: CD4 T cell responses are larger than we think. J Immunol 187, 5568–5576.

Montes de Oca, M., Kumar, R., de Labastida Rivera, F., Amante, F.H., Sheel, M., Faleiro, R.J., Bunn, P.T., Best, S.E., Beattie, L., Ng, S.S., et al. (2016). Blimp-1-Dependent IL-10 Production by Tr1 Cells Regulates TNF-Mediated Tissue Pathology. PLoS Pathog 12, e1005398.

Moormann, A.M., Sumba, P.O., Chelimo, K., Fang, H., Tisch, D.J., Dent, A.E., John, C.C., Long, C.A., Vulule, J., and Kazura, J.W. (2013). Humoral and cellular immunity to Plasmodium falciparum merozoite surface protein 1 and protection from infection with blood-stage parasites. J Infect Dis 208, 149–158.

Nakayamada, S., Kanno, Y., Takahashi, H., Jankovic, D., Lu, K.T., Johnson, T.A., Sun, H.W., Vahedi, G., Hakim, O., Handon, R., et al. (2011). Early Th1 cell differentiation is marked by a Tfh cell-like transition. Immunity 35, 919–931.

Nurieva, R.I., Chung, Y., Hwang, D., Yang, X.O., Kang, H.S., Ma, L., Wang, Y.H., Watowich, S.S., Jetten, A.M., Tian, Q., et al. (2008). Generation of T follicular helper cells is mediated by interleukin-21 but independent of T helper 1, 2, or 17 cell lineages. Immunity 29, 138–149.

Nurieva, R.I., Chung, Y., Martinez, G.J., Yang, X.O., Tanaka, S., Matskevitch, T.D., Wang, Y.H., and Dong, C. (2009). Bcl6 mediates the development of T follicular helper cells. Science 325, 1001–1005.

O’Shea, J.J., and Paul, W.E. (2010). Mechanisms underlying lineage commitment and plasticity of helper CD4+ T cells. Science 327, 1098–1102.

Oakley, M.S., Sahu, B.R., Lotspeich-Cole, L., Solanki, N.R., Majam, V., Pham, P.T., Banerjee, R., Kozakai, Y., Derrick, S.C., Kumar, S., et al. (2013). The transcription factor T-bet regulates parasitemia and promotes pathogenesis during Plasmodium berghei ANKA murine malaria. J Immunol 191, 4699–4708.

Obeng-Adjei, N., Portugal, S., Tran, T.M., Yazew, T.B., Skinner, J., Li, S., Jain, A., Felgner, P.L., Doumbo, O.K., Kayentao, K., et al. (2015). Circulating Th1-Cell-type Tfh Cells that Exhibit Impaired B Cell Help Are Preferentially Activated during Acute Malaria in Children. Cell Rep 13, 425–439.

Oestreich, K.J., Huang, A.C., and Weinmann, A.S. (2011). The lineage-defining factors T-bet and Bcl-6 collaborate to regulate Th1 gene expression patterns. J Exp Med 208, 1001–1013.

Oestreich, K.J., Mohn, S.E., and Weinmann, A.S. (2012). Molecular mechanisms that control the expression and activity of Bcl-6 in TH1 cells to regulate flexibility with a TFH-like gene profile. Nat Immunol 13, 405–411.

Opata, M.M., Carpio, V.H., Ibitokou, S.A., Dillon, B.E., Obiero, J.M., and Stephens, R. (2015). Early effector cells survive the contraction phase in malaria infection and generate both central and effector memory T cells. J Immunol 194, 5346–5354.

Paivandy, A., Calounova, G., Zarnegar, B., Ohrvik, H., Melo, F.R., and Pejler, G. (2014). Mefloquine, an anti-malaria agent, causes reactive oxygen species-dependent cell death in mast cells via a secretory granule-mediated pathway. Pharmacol Res Perspect 2, e00066.

Parish, I.A., Marshall, H.D., Staron, M.M., Lang, P.A., Brustle, A., Chen, J.H., Cui, W., Tsui, Y.C., Perry, C., Laidlaw, B.J., et al. (2014). Chronic viral infection promotes sustained Th1-derived immunoregulatory IL-10 via BLIMP-1. J Clin Invest 124, 3455–3468.

Perez-Mazliah, D., and Langhorne, J. (2014). CD4 T-cell subsets in malaria: TH1/TH2 revisited. Front Immunol 5, 671.

Perez-Mazliah, D., Ng, D.H., Freitas do Rosario, A.P., McLaughlin, S., Mastelic-Gavillet, B., Sodenkamp, J., Kushinga, G., and Langhorne, J. (2015). Disruption of IL-21 signaling affects T cell-B cell interactions and abrogates protective humoral immunity to malaria. PLoS Pathog 11, e1004715.

Perez-Mazliah, D., Nguyen, M.P., Hosking, C., McLaughlin, S., Lewis, M.D., Tumwine, I., Levy, P., and Langhorne, J. (2017). Follicular Helper T Cells are Essential for the Elimination of Plasmodium Infection. EBioMedicine 24, 216–230.

Ray, J.P., Marshall, H.D., Laidlaw, B.J., Staron, M.M., Kaech, S.M., and Craft, J. (2014). Transcription factor STAT3 and type I interferons are corepressive insulators for differentiation of follicular helper and T helper 1 cells. Immunity 40, 367–377.

Roetynck, S., Olotu, A., Simam, J., Marsh, K., Stockinger, B., Urban, B., and Langhorne, J. (2013). Phenotypic and functional profiling of CD4 T cell compartment in distinct populations of healthy adults with different antigenic exposure. PLoS One 8, e55195.

Ryg-Cornejo, V., Ioannidis, L.J., Ly, A., Chiu, C.Y., Tellier, J., Hill, D.L., Preston, S.P., Pellegrini, M., Yu, D., Nutt, S.L., et al. (2016). Severe Malaria Infections Impair Germinal Center Responses by Inhibiting T Follicular Helper Cell Differentiation. Cell Rep 14, 68–81.

Saraiva, M., Christensen, J.R., Veldhoen, M., Murphy, T.L., Murphy, K.M., and O’Garra, A. (2009). Interleukin-10 production by Th1 cells requires interleukin-12-induced STAT4 transcription factor and ERK MAP kinase activation by high antigen dose. Immunity 31, 209–219.

Schulz, E.G., Mariani, L., Radbruch, A., and Hofer, T. (2009). Sequential polarization and imprinting of type 1 T helper lymphocytes by interferon-gamma and interleukin-12. Immunity 30, 673–683.

Stephens, R., Albano, F.R., Quin, S., Pascal, B.J., Harrison, V., Stockinger, B., Kioussis, D., Weltzien, H.U., and Langhorne, J. (2005). Malaria-specific transgenic CD4(+) T cells protect immunodeficient mice from lethal infection and demonstrate requirement for a protective threshold of antibody production for parasite clearance. Blood 106, 1676–1684.

Stephens, R., and Langhorne, J. (2010). Effector memory Th1 CD4 T cells are maintained in a mouse model of chronic malaria. PLoS Pathog 6, e1001208.

Stevenson, M.M., Tam, M.F., Wolf, S.F., and Sher, A. (1995). IL-12-induced protection against blood-stage Plasmodium chabaudi AS requires IFN-gamma and TNF-alpha and occurs via a nitric oxide-dependent mechanism. J Immunol 155, 2545–2556.

Stumhofer, J.S., Silver, J.S., Laurence, A., Porrett, P.M., Harris, T.H., Turka, L.A., Ernst, M., Saris, C.J., O’Shea, J.J., and Hunter, C.A. (2007). Interleukins 27 and 6 induce STAT3-mediated T cell production of interleukin 10. Nat Immunol 8, 1363–1371.

Su, Z., and Stevenson, M.M. (2000). Central role of endogenous gamma interferon in protective immunity against blood-stage Plasmodium chabaudi AS infection. Infect Immun 68, 4399–4406.

Su, Z., and Stevenson, M.M. (2002). IL-12 is required for antibody-mediated protective immunity against blood-stage Plasmodium chabaudi AS malaria infection in mice. J Immunol 168, 1348–1355.

Szabo, S.J., Kim, S.T., Costa, G.L., Zhang, X., Fathman, C.G., and Glimcher, L.H. (2000). A novel transcription factor, T-bet, directs Th1 lineage commitment. Cell 100, 655–669.

Tubo, N.J., and Jenkins, M.K. (2014). TCR signal quantity and quality in CD4+ T cell differentiation. Trends Immunol 35, 591–596.

Tubo, N.J., Pagan, A.J., Taylor, J.J., Nelson, R.W., Linehan, J.L., Ertelt, J.M., Huseby, E.S., Way, S.S., and Jenkins, M.K. (2013). Single naive CD4+ T cells from a diverse repertoire produce different effector cell types during infection. Cell 153, 785–796.

Weinmann, A.S. (2014). Regulatory mechanisms that control T-follicular helper and T-helper 1 cell flexibility. Immunol Cell Biol 92, 34–39.

Weinstein, J.S., Laidlaw, B.J., Lu, Y., Wang, J.K., Schulz, V.P., Li, N., Herman, E.I., Kaech, S.M., Gallagher, P.G., and Craft, J. (2018). STAT4 and T-bet control follicular helper T cell development in viral infections. J Exp Med 215, 337–355.

Wikenheiser, D.J., Brown, S.L., Lee, J., and Stumhofer, J.S. (2018). NK1.1 Expression Defines a Population of CD4(+) Effector T Cells Displaying Th1 and Tfh Cell Properties That Support Early Antibody Production During Plasmodium yoelii Infection. Front Immunol 9, 2277.

Wikenheiser, D.J., Ghosh, D., Kennedy, B., and Stumhofer, J.S. (2016). The Costimulatory Molecule ICOS Regulates Host Th1 and Follicular Th Cell Differentiation in Response to Plasmodium chabaudi chabaudi AS Infection. J Immunol 196, 778–791.

Wu, H., Xu, L.L., Teuscher, P., Liu, H., Kaplan, M.H., and Dent, A.L. (2015). An Inhibitory Role for the Transcription Factor Stat3 in Controlling IL-4 and Bcl6 Expression in Follicular Helper T Cells. J Immunol 195, 2080–2089.

Xu, H., Hodder, A.N., Yan, H., Crewther, P.E., Anders, R.F., and Good, M.F. (2000). CD4+ T cells acting independently of antibody contribute to protective immunity to Plasmodium chabaudi infection after apical membrane antigen 1 immunization. J Immunol 165, 389–396.

Zander, R.A., Vijay, R., Pack, A.D., Guthmiller, J.J., Graham, A.C., Lindner, S.E., Vaughan, A.M., Kappe, S.H.I., and Butler, N.S. (2017). Th1-like Plasmodium-Specific Memory CD4(+) T Cells Support Humoral Immunity. Cell Rep 21, 1839–1852.

